# Structure of cytoplasmic RNA polymerase II

**DOI:** 10.64898/2025.12.10.692585

**Authors:** Annamaria Hlavata, Benjamin Neuditschko, Ulla Schellhaas, Clemens Plaschka, Franz Herzog, Carrie Bernecky

**Affiliations:** Institute of Science and Technology Austria (ISTA), Am Campus 1, 3400 Klosterneuburg, Austria; Institute Krems Bioanalytics, IMC University of Applied Sciences, 3500, Krems, Austria; Research Institute of Molecular Pathology (IMP), Vienna BioCenter (VBC), Campus Vienna-Biocenter 1, 1030, Vienna, Austria

## Abstract

RNA polymerase II (Pol II) must be assembled in the cytoplasm before it enters the nucleus, where it transcribes protein-coding genes. Although transcription by Pol II is intensively studied, how this central multi-subunit enzyme is made and the role of dedicated factors remains unclear. Here, we report the integrative structural analysis of a native human Pol II from the cytoplasm captured near the end of biogenesis. The complex contained Gdown1 and three biogenesis factors – RPAP2 and the critical small GTPases GPN1 and GPN3. Cryo-EM analysis of the complex revealed how Gdown1 and RPAP2 associate with Pol II and prevent the premature association of transcription factors. Further biochemical and cryo-EM analysis revealed how RPAP2 recruits GPN1–GPN3 to the complex, and how the assembly of the RPAP2–GPN1–GPN3 complex is controlled by GTP hydrolysis. The combined results uncover a network of interactions that chaperone cytoplasmic Pol II to prevent aberrant interactions, reveal a GTP-controlled switch during the final stages of Pol II biogenesis, and suggest a general mechanism for the action of GPN-loop GTPase family of enzymes.

## Introduction

RNA polymerase II (Pol II) is the eukaryotic enzyme responsible for mRNA production. Although Pol II carries out transcription in the nucleus, it must first be made in the cytoplasm where it is assembled from its 12 constituent subunits, before it is transported into the nucleus as a fully-formed complex^1^. The assembly of Pol II has been shown to require the action of a variety of biogenesis factors, including chaperones^1^, RNA polymerase-associated proteins (RPAPs)^2^, and small GPN-loop GTPases (GPNs)^3,4^. However, the roles of these factors and the mechanisms involved have remained unclear. Regulation of Pol II assembly and nuclear import is important for providing sufficient enzyme to daughter cells after cell division, as well as to cells recovering from DNA damage, during which Pol II is degraded^5^. Furthermore, changes in levels of Pol II have been shown to control gene regulation^5–8^. Taken together, Pol II assembly is critical for gene expression and its regulation.

Insights into the factors participating in assembly and import of Pol II have come from affinity purification of Pol II- and Pol II-subcomplex-associated factors from soluble extracts, as well as from functional studies examining the effects of depletion or knockout^9^. At steady state, soluble Pol II interacts with several factors including chaperones, RNA-polymerase associating proteins (RPAPs), GPN-loop GTPases (GPNs), and the metazoan-specific Gdown1^3,10,11^. Experiments performed after the induction of Pol II degradation have revealed the existence of partially assembled Pol II complexes that are pre-associated with several factors including GPNs, RPAPs, Gdown1, and chaperones. Thus, GPN and RPAP proteins associate with Pol II subunits before the assembly of the complete enzyme, potentially participating in the merging of these subcomplexes and the incorporation of RPB5^1^. The assembled cytoplasmic Pol II has also been shown to associate with RPAP2 and GPN1-GPN3^4^, and Gdown1 is known to very stably bind to 12-subunit Pol II^12^. RPAP2, GPN1-3, and Gdown1 are each essential or important for cell growth and development, as evidenced by CRISPR screens in cancer cell lines^13,14^ and knockout and depletion experiments^2–4,15^.

Prior work has indicated important roles for RPAP2 and GPN proteins in Pol II assembly and import. GPN1 has been shown to form a heterodimer with GPN2 or GPN3, with GPN2 implicated in early steps of assembly whereas GPN3 appears to function later^16,17^. The GPN1-GPN3 complex is the predominant form of GPN1 in mammalian cells^18^, and depletion of either protein or mutation of the GTP-binding pocket of GPN1 reduces levels of nuclear Pol II^4^. Similarly, loss of RPAP2 leads to defects in Pol II assembly or nuclear import. GPN1 and the C-terminus of RPAP2 have been reported to interact and interestingly, GPN1 is required for the nuclear export and cytoplasmic recycling of RPAP2^2^.

Gdown1^19–21^ (also POLR2M, GRINL1A) has also been identified as an interactor of Pol II and Pol II subcomplexes, and may play a role in Pol II assembly^1^. Gdown1 contains two nuclear export sequences (NES’s) and a cytoplasmic anchoring sequence (CAS)^19^, which keeps Gdown1 predominantly localized to the cytoplasm except for during transcriptionally silent developmental stages^15^, stress^19^, and mitosis^20^, during which Gdown1 is phosphorylated, particularly at S270^22^. A plethora of *in vitro* biochemical data indicates that Gdown interferes with all stages of transcription (including initiation, elongation, and termination), but acute Gdown1 depletion has minimal effects on transcription^20^. Altogether, the data indicate a role for Gdown1 during mitosis and during the cellular stress response, likely helping to shut down transcription. However, the interplay between newly assembled Pol II, biogenesis factors, and Gdown1 remains unclear.

Here, we obtain molecular insights into Pol II assembly and how Pol II is chaperoned in the cytoplasm prior to its import into the nucleus. We apply integrative structural analysis of endogenous and *in vitro* reconstituted complexes accompanied by biochemical assays, revealing how the essential factors Gdown1, RPAP2, GPN1, GPN3, and a GTP-gated switch may control the final stages of cytoplasmic Pol II biogenesis.

## Results

### Identification of the Gdown1-associated proteome in cytoplasmic and nuclear fractions

To investigate late events in human Pol II biogenesis, we set out to purify cytoplasmic Pol II complexes. To avoid perturbation of any Pol II surface, we utilized Gdown1 as a handle, given the high stability of the Gdown1-Pol II interaction^22^, Gdown1’s predominantly cytoplasmic localization^15,19,20^, and the significant fraction (30-50%) of Pol II-Gdown1 complexes in whole-cell extracts^12^. We generated a K562 cell line in which both endogenous copies of Gdown1 were tagged with eGFP followed by a 3C protease cleavage site, to allow affinity purification of Gdown1-associated complexes and their elution from the resin under mild conditions (Extended Data Fig. 1a). We then characterized the Gdown1 interactome in both the cytoplasm and nucleoplasm using immunoprecipitation and mass spectrometry analysis, observing many significantly enriched proteins (Fig. 1a,b, Extended Data Fig. 1b-d, and Supplementary Table 1). All subunits of Pol II (RPB1-12) were significantly enriched, consistent with the predominant accumulation of assembled Pol II at steady state. We also identified many proteins previously shown to interact with Pol II, including RPAP1-3, GPN1-3, DSIF (SUPT5H/SUPT4H), RECQL5, TFIIS (TCEA1), and the Mediator and Integrator complexes^1,3,10,11^ (Extended Data Fig. 1c,d).

**Figure 1.**
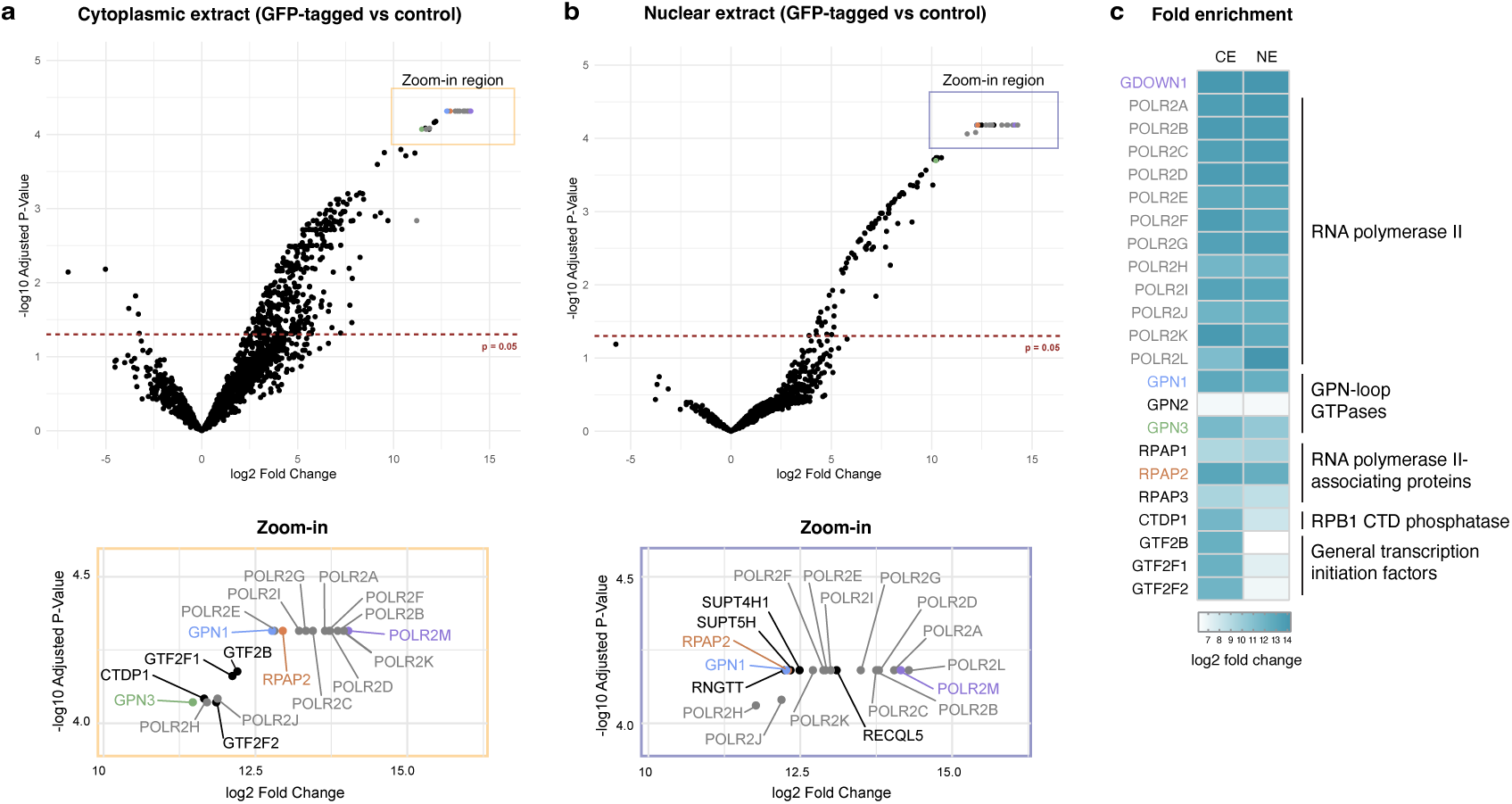
Identification of Gdown1-interacting factors. **a**, Volcano plot (full and zoom-in view) of the proteins identified in MS analysis following IP of Gdown1 from cytoplasmic extract. **b**, Volcano plot (full and zoom-in view) of the proteins identified in MS analysis following IP of Gdown1 from nucleoplasmic extract. **c**, Heatmap comparing the abundance of top cytoplasmic Gdown1-associating proteins, RPAP proteins, and GPN proteins recovered from the cytoplasmic and nuclear fractions, with darker blue color representing increasing fold change.

Comparison of the most significantly enriched cytoplasmic and nucleoplasmic Gdown1 interactors revealed high enrichment of RPAP2, GPN1, and GPN3 in both cytoplasm and nucleoplasm (Fig. 1c). Given the enrichment of known Pol II biogenesis factors, a significant fraction of the nuclear Pol II-Gdown1 complex may represent Pol II that was newly imported into the nucleus, but which has not yet engaged in transcription, and may suggest that Gdown1, RPAP2, GPN1 and GPN3 are imported together to the nucleus from the cytoplasm. Although no structural information regarding GPN1 and GPN3 interaction with Pol II is available, structures of Pol II-RPAP2 have revealed how the N-terminal Pol II-interacting domain of RPAP2 binds to the Pol II cleft and blocks nucleic acid association^23,24^. Altogether, Gdown1 pulldown efficiently recovered a cytoplasmic Pol II complex.

### Architecture of endogenous cytoplasmic RNA polymerase II

The analysis above suggested that a Pol II complex enriched in Gdown1, RPAP2, GPN1, and GPN3 could be isolated from the cytoplasm. To gain insight into the architecture of this complex prior to nuclear import, we aimed to determine its structure. Gdown1-containing complexes were purified from the cytoplasm of K562 cells endogenously expressing GFP-tagged Gdown1, followed by GraFix separation to enrich for Pol II-containing complexes. Cryo-EM analysis of the resulting complexes revealed the structure of Pol II bound to Gdown1 and RPAP2 at a nominal resolution of 2.7 Å, in which 12-subunit Pol II, the Pol II-binding region of RPAP2, and three distinct Pol II-binding regions of Gdown1, but no nucleic acid density, could be resolved (Fig 2a,b, Extended Data Fig. 2). RPAP2 was bound to the RPB1 jaw and RPB5, essentially as previously observed^23,24^, and the Pol II clamp and active site were disordered. However, due to improved density, and a previously existing model for Pol II-Gdown1^15^ could be generated, in which the register could be assigned for the N-terminal regions of Gdown1, the connectivity of two Gdown1 regions could be clarified, and the Gdown1 region binding to the RPB2 protrusion could be extended (Extended Data Fig. 2j). In the new model, a more extensive binding interface with RPB2 could be observed and three Gdown1-Pol II interfaces could be identified, including an RPB10-binding domain (interface 1), an external-binding domain (interface 2), and a protrusion-binding domain (interface 3). Two nuclear export sequences and one cytoplasmic anchoring sequence are localized in unstructured regions, and thus likely to be functional in the context of the Pol II-Gdown1 complex. Phosphorylation of Gdown1 at serine 270 has been shown to occur during mitosis and reduce the affinity of Gdown1 for Pol II early elongation complexes^22^. This residue of Gdown1 is located in a disordered region, suggesting that this residue is available for phosphorylation in the context of the Pol II-Gdown1 complex, and that additional protein or nucleic acid factors may be required to decrease affinity to Pol II. To validate our model, we performed chemical crosslinking combined with mass spectrometry (XL-MS) of the cytoplasmic sample. Although few inter-subunit crosslinks were observed, all were consistent with the Pol II-Gdown1-RPAP2 structure and previous XL-MS analysis of a Pol II-Gdown1 binary complex^15^, suggesting that the Pol II-Gdown1 interface is the same in the presence of additional factors present in the cytoplasm (Extended Data Fig. 3a,b).

**Figure 2.**
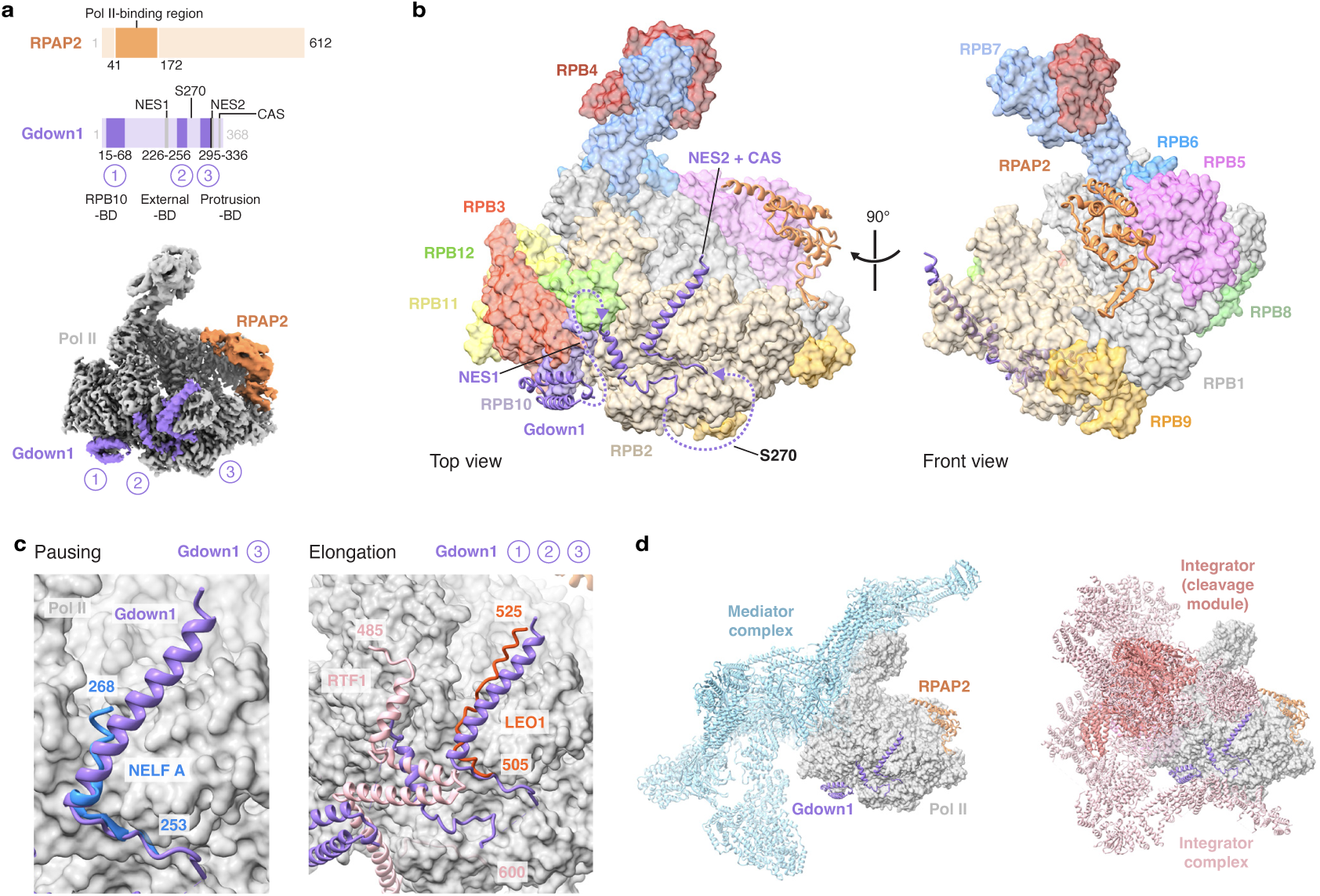
Structural analysis of cytoplasmic Pol II. **a**, Top, schematic of RPAP2 and Gdown1, with regions resolved in the cytoplasmic Pol II structure shown in dark orange and purple, respectively. The Gdown1 RPB10-binding domain (interface 1), external-binding domain (interface 2), and protrusion-binding domain (interface 3) are indicated. Bottom, top view of the composite cytoplasmic Pol II reconstruction. Pol II, gray; RPAP2, orange, Gdown1, purple. **b**, Top and front view of the Pol II-Gdown1-RPAP2 structure, with Pol II colored by subunit. **c**, Superposition of the Pol II-Gdown1-RPAP2 structure with a structure of paused Pol II bound to DSIF and NELF (left, PDB ID 8UHD^72^) and a structure of an activated Pol II PAF1C and RTF1-containing elongation complex (right, PDB ID 6TED^73^). The Gdown1 regions involved in clashes are noted with circled numbers. Colored as follows: NELF A, blue; LEO1, red; RTF1, light pink. **d**, Comparison of the Pol II binding interfaces of the Mediator (left, PDB ID 7ENA^31^) and Integrator (right, PDB 8RBX^28^) complexes with those of Gdown1 and RPAP2. Colored as follows: Mediator, light blue; Integrator, pink; Integrator cleavage module, dark pink.

Gdown1 has been shown *in vitro* to inhibit all stages of transcription, including initiation^12^, promoter-proximal pausing^25^, elongation^20^, and termination^22^. Structural and biochemical competition between Gdown1 and TFIIF has been well characterized, and as expected, we observed a structural clash between the Gdown1 protrusion-binding domain and the small subunit of TFIIF, as well as a small clash with TFIIB. This structural incompatibility, together with biochemical disruption of the Pol II-TFIIF interaction by RPAP2^24^, suggests that the cytoplasmic enrichment of TFIIF (GTF2F1, GTF2F2) in the cytoplasmic Gdown1 IP-MS is due to an alternative protein-protein interaction (Fig. 1a, Supplementary Text 1). Comparison of the Pol II-Gdown1-RPAP2 structure to an AlphaFold3-predicted structure of Pol II and the mitotic transcription termination factor TTF2 revealed that the Gdown1 RBP10-binding domain is responsible for the observed biochemical competition between the factors (Extended Data Fig. 3c). Gdown1 has also been shown to relieve repression by the pausing factor NELF^25^. This effect can be explained by a structural clash between the Gdown1 protrusion-binding domain and the NELF A subunit of the NELF complex (Fig. 2c). Furthermore, previously observed functional competition between Gdown1 and the PAF1 complex (PAF1C) with RTF1^20^ can be explained by a clash between the PAF1C subunit LEO1 and the protrusion-binding domain of Gdown1 and between RTF1 and the RBP10- and external-binding domains of Gdown1 (Fig. 2c).

Notably, Gdown1 does not appear to interfere with the binding of many essential transcription factors. For example, comparison to a structure of Mediator within a TFIID-containing pre-initiation complex revealed no direct competition between Mediator and Gdown1 (Fig. 2d). Thus, the previously reported activity of the Mediator complex in relieving Gdown1-based transcriptional repression^12^ is likely due to Mediator-dependent stimulation of pre-initiation complex assembly, for example through stabilization of TFIIB binding^26^, allowing general transcription initiation factors to displace Gdown1 binding to their interfaces. Structural compatibility is also observed between Gdown1 and the transcription elongation factor DSIF, a platform for the binding of many transcriptional cofactors and which associates with Pol II shortly after promoter escape until transcription termination^27^, consistent with previous biochemical data^25^ (Extended Data Fig. 3d). Furthermore, Gdown1 binding is also compatible with association of the Integrator complex, despite the extensive binding interfaces between Integrator and Pol II^28,29^ (Fig. 2d, Extended Data Fig. 3e). Integrator is a large multi-subunit complex with RNA cleavage activity and key roles in termination of noncoding RNAs and promoter-proximally paused Pol II^30^. Thus, Gdown1 would be located in a way consistent with displacement of the pausing factor NELF without impeding the pausing termination activity of Integrator. Inspection of enriched proteins in the Gdown1 IP-MS experiments revealed that Mediator, DSIF, and the Integrator complex were all enriched, particularly in the pulldown from nucleoplasmic extracts, consistent with the interactions of Mediator and Integrator with the CTD of the largest subunit of Pol II^31,32^ and the structural compatibility between Gdown1 and these complexes when bound to Pol II (Extended Data Fig. 1c,d).

### Analysis of a reconstituted RNA polymerase II-Gdown1-RPAP2-GPN1-GPN3 complex

Analysis of the proteins that co-migrated with Pol II in sucrose gradient centrifugation (Extended Data Fig. 4a), as well as a body of previously published work^4,10^ suggests that Pol II, RPAP2, Gdown1, and GPN1-3 exist as part of a single complex in the cytoplasm. Additionally, previous work has indicated a direct interaction between GPN1 and both Pol II and RPAP2 to GPN1^2^. However, cryo-EM and XL-MS analysis of the endogenous cytoplasmic Pol II complex did not reveal the location of GPN1 and GPN3, despite their relatively high enrichment in IP-MS in both cytoplasmic and nuclear Gdown1 complexes. To investigate the interactions of GPN1-GPN3 within the cytoplasmic Pol II complex, we reconstituted a complex containing Pol II, Gdown1, RPAP2, and GPN1-GPN3 from purified components (Extended Data Fig. 4b,c). We assessed complex formation by mass photometry and size-exclusion chromatography, both of which indicated assembly of the Pol II–Gdown1–RPAP2–GPN1–GPN3 complex (Fig. 3a,b). Mass photometry was applied to independently assess the interactions of Pol II with RPAP2 and GPN1-GPN3. Isolated Pol II was primarily dimeric, in line with previous work showing that purified Pol II forms a dimeric complex, which has been suggested to be a storage form of Pol II^33,34^. Recombinant RPAP2 readily bound the Pol II complex, incorporating into both Pol II monomeric and dimeric forms, but decreasing the fraction of Pol II dimers. Separately, recombinant GPN1-GPN3 shifted a small fraction of monomeric Pol II, but did not incorporate into the dimer, suggesting a direct but relatively weak Pol II-GPN interaction. When added together, the presence of Pol II dimer was strongly reduced, and Pol II was quantitatively shifted into a monomeric Pol II-RPAP2-GPN1-GPN3 complex, indicating that RPAP2 is required to assemble a stable complex with GPN1-GPN3. Gdown1, which has been previously shown to reduce Pol II dimer formation on its own^34^, readily incorporated into this monomeric Pol II-RPAP2-GPN1-GPN3 complex (Fig. 2b). Taken together, these observations indicate that cytoplasmic Pol II exists predominantly in a monomeric form, consistent with the lack of observed dimers in cryo-EM of the native, cytoplasmic Pol II.

**Figure 3.**
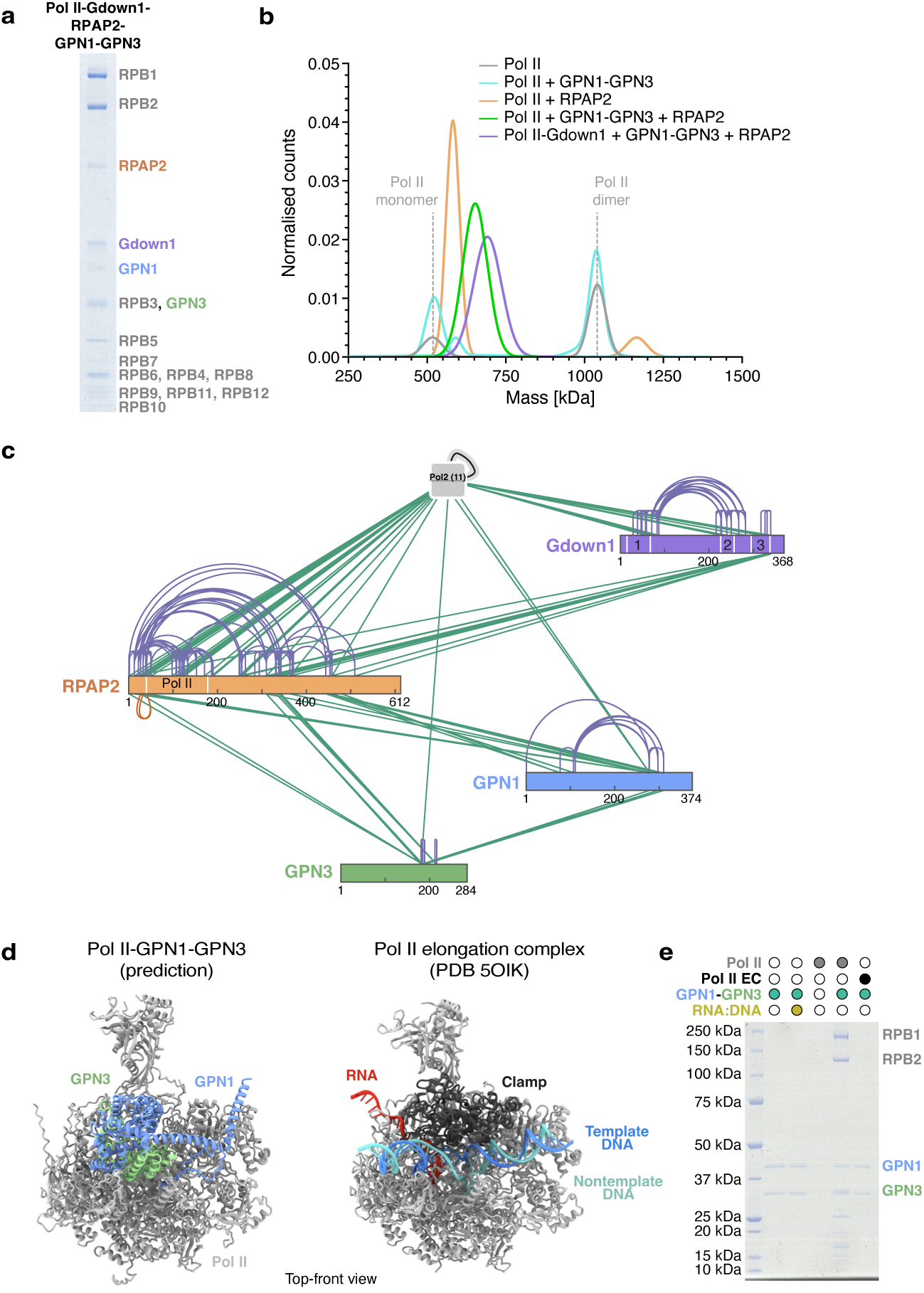
Architecture of the Pol II-Gdown1-RPAP2-GPN1-GPN3 complex. **a**, SDS-PAGE analysis (Coomassie) of size-exclusion-purified recombinant Pol II-Gdown1-RPAP2-GPN1-GPN3. **b**, Mass photometry analysis of Pol II alone or in the presence of RPAP2, GPN1-GPN3, and/or Gdown1. Molecular weights corresponding to the unbound Pol II monomer and dimer are indicated with dashed gray lines. **c**, Crosslinking-mass spectrometry of recombinant Pol II-Gdown1-RPAP2-GPN1-GPN3. Violet, self crosslinked peptides; green, heteromeric crosslinked peptides. Generated by xiView. **d**, Top-front views of the Pol II-GPN1-GPN3 structure prediction and the Pol II elongation complex (Pol II EC, PDB ID 5OIK^74^). Colored as follows: Pol II, gray; GPN1, blue; GPN3, green; Pol II EC clamp, black; RNA, red; template DNA, blue, non-template DNA, cyan. **e,** Pull-down of Pol II and Pol II elongation complex (Pol II EC) via immobilized His-GPN1-GPN3.

To gain further insights into the architecture of the complex, we identified chemical crosslinks of the reconstituted Pol II-Gdown1-RPAP2-GPN1-GPN3 by mass spectrometry. The obtained crosslinks were consistent with the cytoplasmic Pol II structure (Fig. 3c, Extended Data Fig. 4d,e). We observed a large number of inter-protein crosslinks, including multiple links between the unstructured N-terminal region of RPAP2 and Pol II, which were consistent with a localization near the Pol II clamp and DNA binding cleft. This region also crosslinked to the Gdown1 protrusion-binding domain (interface 3), consistent with a localization near the Pol II DNA-binding cleft, and to GPN1 and GPN3. We observed multiple crosslinks between GPN1-GPN3 and the C-terminus of RPAP2, consistent with previous observations that the C-terminus of RPAP2 is required for its association with GPN1^2^. Furthermore, GPN1-GPN3 crosslinked to RPB1, the largest subunit of Pol II, through its clamp and C-terminal domains(Extended Data Fig. 4f). Structure prediction of Pol II-GPN1-GPN3 suggested that GPN1-GPN3 interacts with the DNA-binding cleft of Pol II through electrostatic interactions with negatively charged regions of GPN1 and especially GPN3, with the negatively charged GPN3 C-terminal tail predicted to associate with the pore and tunnel which accommodate backtracked RNA in the arrested elongation complex^35^ (Fig. 3d, Extended Data Fig. 4g,h). This interaction would require an open clamp conformation, and would be blocked in a Pol II elongation complex. To test this, we incubated GPN1–GPN3 with apo or nucleic acid-bound forms of Pol II in a pulldown binding assay (Fig. 3e). Whereas GPN1–GPN3 was able to associate with Pol II, it did not interact with the Pol II elongation complex, suggesting that its primary binding site lies within the DNA-binding cleft or clamp domain rather than the stalk (RPB4/7) or the C-terminal domain (CTD) as previously suggested^4^. Altogether, these results suggest that GPN1-GPN3 interacts directly and likely transiently with the Pol II DNA binding cleft, and is anchored to the Pol II complex through the flexibly associated, unresolved C-terminal region of RPAP2.

### Structural and biochemical analysis of RPAP2-GPN1-GPN3

To visualize how GPN1-GPN3 is anchored to the cytoplasmic Pol II complex through RPAP2, we reconstituted a RPAP2-GPN1-GPN3 complex. This complex was readily observed in mass photometry analysis and size exclusion chromatography (Fig. 4a,b). Pulldown experiments utilizing purified proteins showed that the Pol II-binding region of RPAP2 is not able to directly interact with GPN1-GPN3 (Extended Data Fig. 5a), suggesting that the observed crosslinks between the RPAP2 N-terminus and GPN1-GPN3 were due to proximity in the context of the Pol II-Gdown1-RPAP2-GPN1-GPN3 complex, rather than a direct interaction between the two. Cryo-EM analysis of the purified complex resulted in a 2.8-Å nominal resolution structure, into which the GPN1-GPN3 heterodimer and a C-terminal region of RPAP2 could be modeled (Fig. 4c-e, Extended Data Fig. 5b-f). GPN1-GPN3 formed the core of the structure, with the very C-terminal ends of both flexible and solvent-exposed, including the GPN1 nuclear export signal. RPAP2 formed an extended interface with the GPN1-GPN3 dimer. A largely alpha-helical, GPN1-binding domain of RPAP2 was formed by residues 359-389 and 490-608, associating with GPN1 in close proximity to the GTP/GDP binding pocket. A second region of RPAP2 encompassing residues 428-467 bound to the GPN1-GPN3 dimerization interface, continuing along GPN1. A hydrophobic patch previously suggested to associate with disordered hydrophobic peptides^36^, was buried within the GPN1-GPN3 complex (Extended Data Fig. 5g). However, several surface-exposed hydrophobic residues are located at the interface between GPN1 and GPN3, suggesting that other binding pockets may allow allosteric activation of GPN1 GTPase activity as previously described^36^ (Extended Data Fig. 5h). Mapping the crosslinks of the Pol II-RPAP2-GPN1-GPN3 complex onto the structure RPAP2-GPN1-GPN3 showed overall good agreement with the experimentally determined model compared with an AlphaFold3-predicted model (Extended Data Fig. 5i,j), validating the interaction of RPAP2 with GPN1 in proximity to the GPN3 nucleotide-binding pocket.

**Figure 4.**
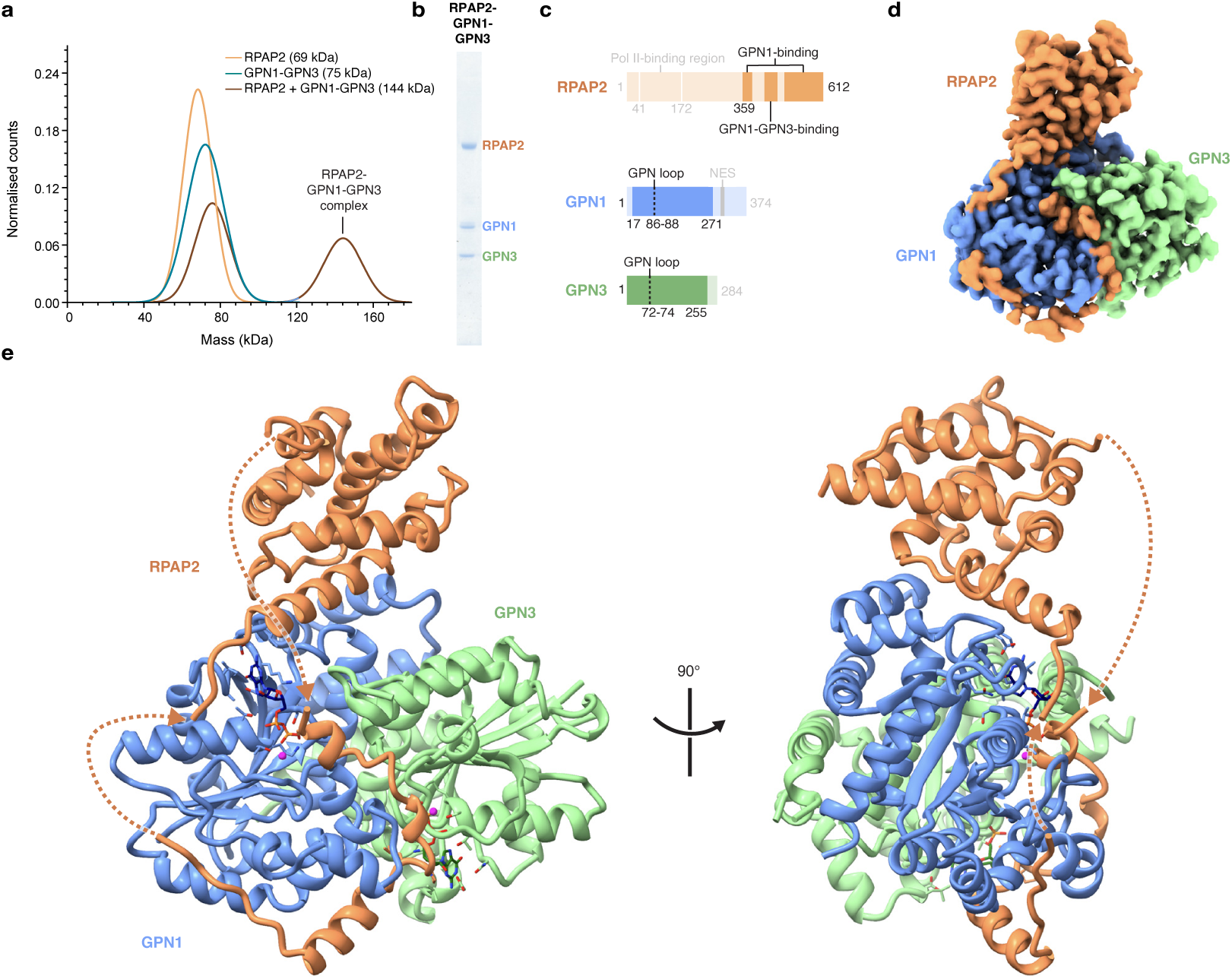
Structural analysis of RPAP2-GPN1-GPN3. **a**, Mass photometry analysis of RPAP2, GPN1-GPN3, and RPAP2-GPN1-GPN3. **b**, SDS-PAGE analysis (Coomassie) of size-exclusion-purified RPAP2-GPN1-GPN3. **c,** Schematic of RPAP2, GPN1, and GPN3 proteins, in which unresolved regions of the RPAP2-GPN1-GPN3 structure of at least 10 amino acids in length are shown in faded colors. RPAP2, orange; GPN1, blue; GPN3, green. **d,** Cryo-EM density map of RPAP2-GPN1-GPN3 colored as in (c). **e,** Two views of the RPAP2-GPN1-GPN3 structure, with the connectivity of the unresolved regions of RPAP2 indicated with orange dashed lines.

Inspection of the density within the nucleotide binding pockets of GPN1 and GPN3 revealed that both were bound to a molecule of GDP, which had been co-purified with the complex after recombinant expression in insect cells (Extended Data Fig. 5k). This observation coupled to the conservation of the G1-G3 and GPN motifs^36^ indicates that in the context of a GPN1-GPN3 heterodimer, both GPN1 and GPN3 are capable of GTP hydrolysis. The GPN-loops of both GPN1 and GPN3 were positioned away from the bound nucleotide, consistent with a GDP-bound state (Extended Data Fig. 5l). RPAP2 passed directly next to the GDP molecules of both GPN1 and GPN3, with a mode of interaction is incompatible with a GTP-bound state (Fig. 5a-c). This observation is in agreement with previous work suggesting that GDP may increase the RPAP2-GPN1 binary interaction^2^. In the presence of GTP or its non-hydrolyzable analogue, the GPN loop is involved in the coordination of the γ-phosphate, leading to a more closed dimeric arrangement^37^. These data suggested that RPAP2 recognizes the GDP-bound conformation of the GPN1-GPN3 complex. To test this, we performed mass photometry measurements of GPN1-GPN3 and RPAP2 complex formation in the presence of either no additional nucleotide, GDP, GTP, or the non-hydrolyzable GTP analogue GppNHp. Whereas the complex readily formed without additional nucleotide or in the presence of GDP, RPAP2 binding was inhibited in the presence of non-hydrolyzable GTP, and even further reduced in the presence of GTP (Fig. 5d,e). Complementary pulldown experiments qualitatively confirmed the reduction in RPAP2 affinity for GPN1-GPN3 in the presence of GTP (Extended Data Fig. 5m). Together, these results identify RPAP2 as a sensor of GPN1-GPN3 state and allow modeling of a cytoplasmic Pol II structure in which GPN1-GPN3 are anchored to the complex through RPAP2.

**Figure 5.**
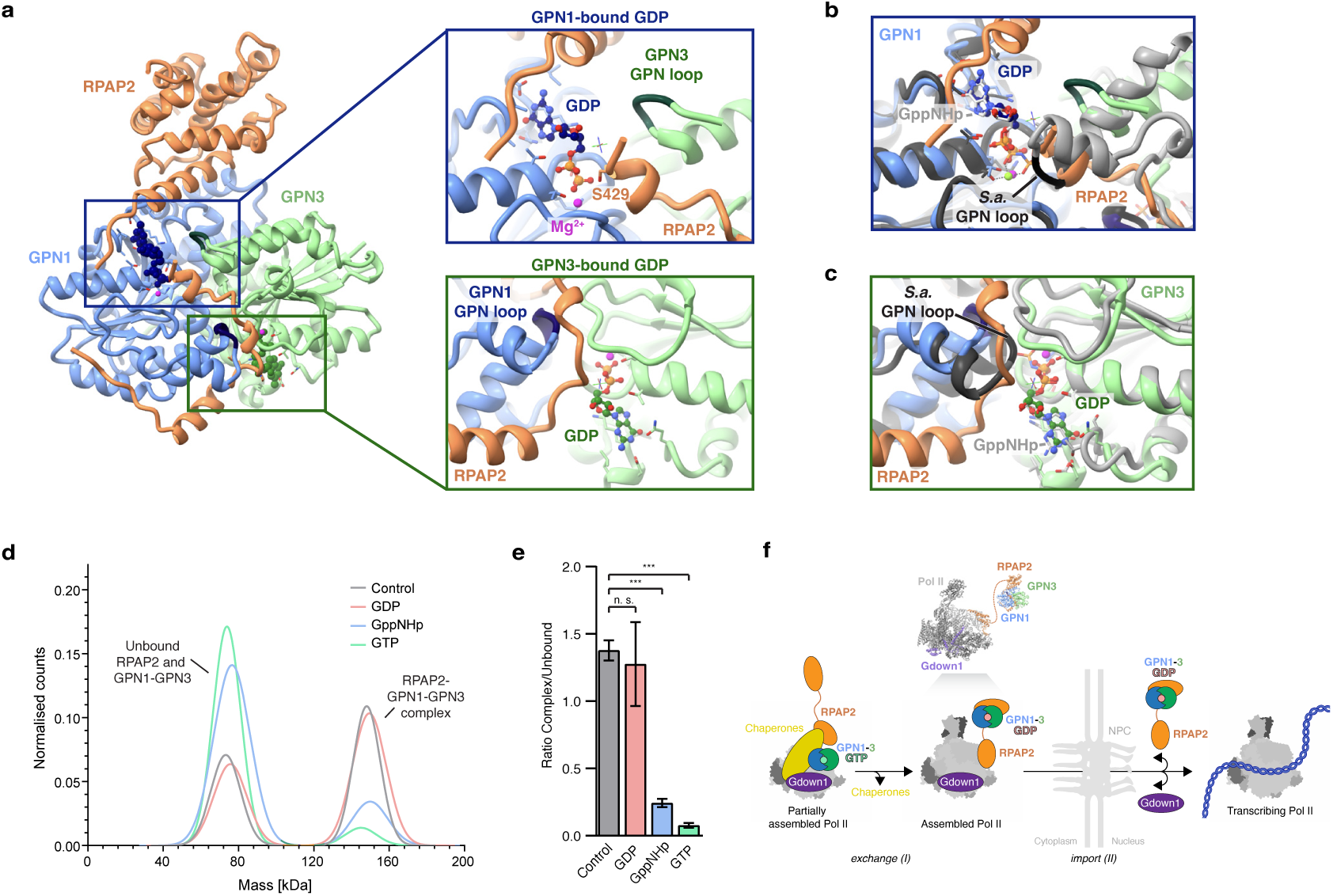
Biochemical and structural analysis of GPN1-GPN3 bound to RPAP2. **a**, A view of the GPN1 and GPN3 nucleotide-binding sites with bound GDP. Left, overview image; right, zoom-in views of the GPN1 (top) and GPN3 (bottom) binding pockets. GPN3 GPN loop, dark green; GPN1 GPN loop, dark blue; Mg^2+^ ions, magenta. **b**-**c**, Superposition of the human RPAP2-GPN1-GPN3 structure (this study) with that of the archeal SaGPN dimer bound to the non-hydrolyzable GTP analogue GppNHp (PDB ID 7ZHF^37^**b,** View of GPN1-bound GDP. Colored as follows: SaGPN monomer 1, dark gray; SaGPN monomer 2, gray; SaGPN-bound GppNHp, light gray, SaGPN-loop black; RPAP2-GPN1-GPN3 structure as in (a). **c**, View of GPN3-bound GDP. Colored as in (b). **d**, Mass-photometry of RPAP2-GPN1-GPN3 complex formation after GPN1-GPN3 pre-treatment with GTP, GppNHp, or GDP. **e**, Ratio of the counts of RPAP2-GPN1-GPN3 complex versus unbound proteins as in (d), performed in triplicate. n.s not significant, **** p<0.0001 **f**, final schematic.

## Discussion

The structural and biochemical analysis of cytoplasmic Pol II provides insights into its biogenesis and the functions of key interactors, with implications for Pol II regulation in the nucleus and cytoplasm.

### Gdown1-dependent inhibition of Pol II-factor binding

The revised Gdown1 structure presented here clarifies the interactions responsible for the transcription inhibition activity of Gdown1, a phenomenon first recognized 20 years ago^12^. One intriguing prior observation is that Gdown1 inhibits pausing *in vitro*^25^, which we could explain as an effect of competition with the pausing factor subunit NELF A. At the same time, Gdown1 has been described to localize together with paused Pol II in ChIP experiments^25^. One intriguing possibility is that Gdown1 may be able to incorporate into a stalled Pol II complex on the way to its termination, for example through the action of Integrator.

The association of Gdown1 with cytoplasmic Pol II complexes together with its role in mitosis and response to stress suggests a possibility for crosstalk and coordination between Pol II assembly and cell growth. For example, Gdown1 may help enforce a requirement for Mediator as transcription is re-started following DNA replication and localization of factors is being re-restablished. Further work is required to elucidate the role of Gdown1 in nucleoplasm, especially its role during mitosis and stress response.

### Pol II-Gdown1 interactions in the context of Pol II biogenesis

Gdown1 directly contacts the majority of the Pol II subunits thought to belong to the RPB2 subcomplex assembly intermediate, including RPB2, RPB3, RPB10, and RPB12^1^. Previous mass spectrometry work has suggested that a RPB1 subcomplex also associates with Gdown1 as well as with the R2TP/Prefoldin-like cochaperone complex, which contains many proteins including Unconventional prefoldin RPB5 interactor 1 (URI1)^1^. Structural prediction suggests that Gdown1 and URI1 directly interact in a way that would conflict with the interaction between Gdown1 and the RPB2 protrusion domain (interface 3)^38^. Speculatively, after the RPB1 and RPB2 subcomplexes are merged, Gdown1 may release URI1 in favor of the three interfaces described in this study. Further work, for example Gdown1 interactome analysis under conditions in which Pol II assembly is stimulated, will be required to investigate a role of Gdown1 in Pol II assembly.

### Interaction of Pol II with factors required for assembly and nuclear import

Although GTP binding was required for yeast GPN1 to immunoprecipitate Pol II^39^, this is not the case for mammalian Pol II, as we observed binding between immobilized GPN1-GPN3 (purified in a GDP-bound state) and Pol II in the absence of added nucleotide. This may be due to the flexible clamp domain of human Pol II, which allows GPN1-GPN3 to associate with the DNA/RNA binding cleft even after RPB1 has joined the assembly, whereas in yeast assembly of the RPB1 clamp might block GPN1-3 binding. GTP binding by GPN1 may facilitate interaction with another protein that binds Pol II and thus this observed interaction is indirect. The human Pol II-GPN1-GPN3 structure prediction suggests that GPN1–GPN3 binding would sterically prevent DNA engagement and clamp closure, similar to the reported ability of RPAP2 to compete with downstream DNA^23,24^. Thus, the DNA binding cleft of Pol II would be chaperoned through protein-protein interactions, which may reduce spurious contacts with nucleic acids in the cytoplasm.

### Function and mechanism of GPN-loop GTPases

GPN1, GPN2, and GPN3 are small eukaryotic GTPases that are required for Pol II assembly and import^3,4^. They belong to a class of proteins known as GPN-loop GTPases, named after a glycine-proline-asparagine (GPN) motif required for their catalytic activity^40^. The GPN-loop GTPase family is conserved from archaea to eukaryotes^4,36,37^ and these enzymes function as constitutive homodimers or heterodimers, independent of GTP- or GDP-bound state^18^. Although homodimers have been observed for recombinantly expressed, purified yeast^36^and human^41^ GPN1, in a cellular context, eukaryotic GPN1 forms heterodimers with either GPN2 or GPN3^42^. In eukaryotes, the majority of GPN1 exists within a heterodimeric complex with GPN3^18^. Thus, study of the heterodimeric state is necessary to understand eukaryotic GPN-loop GTPase structure and function.

Despite X-ray crystallographic studies of two archaeal GPN-loop GTPases^37,40^and isolated yeast Gpn1^36^, how GPN-loop GTPases fulfill their function and the role of GTP hydrolysis in this process remains enigmatic. Analysis of the GTP- and GDP-bound states of the GPN enzyme from the archaea *Sulfolobus acidocaldarius* showed that GTP hydrolysis leads to an allosteric change in the dimerization interface, converting between a closed (GTP-bound) conformation and an open (GDP-bound) conformation^37^. Despite this insight, how this subtle conformational change links to GPN function remained unclear. Furthermore, previous structural and biochemical analysis of isolated yeast Gpn1 (Npa3) revealed that the presence of hydrophobic peptides could stimulate the GTP hydrolysis activity of Gpn1^36^. In this work, the structure of GDP-bound GPN1-GPN3 bound to RPAP2 reveals an extended interaction with GPN1 and the GPN1-GPN3 dimerization interface, with the interface interaction requiring GPN1-GPN3 to be in a GDP-bound conformation. The biochemical results presented here suggest that GTP binding, and perhaps more potently the conformational changes undergone during GTP hydrolysis, destabilize RPAP2 binding. Thus, RPAP2 is the first known effector of GPN-loop GTPases, suggesting a general mechanism by which the GPN dimerization interface, specifically the regions adjacent to the nucleotide-binding pockets, serves as a regulated binding site for effectors.

### Expanded model of Pol II biogenesis

These insights suggest a potential mechanism for GPN1-GPN3 function in Pol II biogenesis as a marker and critical molecular switch of the assembly process. GPN1-GPN3 have been found to associate with partially assembled Pol II complexes, particularly a RBP2- and Gdown1-containing subcomplex before the incorporation of RPB1^1^. Thus, GPN1-GPN3 might associate first with Pol II subcomplexes, possibly via interaction with RPB2 or a chaperone-associated factor, such as URI1. URI1 has been not only predicted to bind Gdown1, but also to interact with GPN1 and GPN3^38^. Interaction with early assembly factors, for example URI1, or exposed hydrophobic peptides may act as guanine exchange factors to keep GPN1-GPN3 in a predominantly GTP-hydrolyzing state, and potentially facilitating a chaperone function. Once assembly of the RPB2-subcomplex is complete, GPN1-3 would revert to a GDP-bound state, facilitating high-affinity interaction with RPAP2. RPAP2 could facilitate the joining of the RBP1 subcomplex, and remain bound to assembled Pol II through the DNA-binding cleft. This would be consistent with previous genetic work in yeast suggesting that the yeast RPAP2 homologue Rtr1 cooperates with the yeast GPN1-GPN3 homologues (Npa3, Gpn3) to facilitate the assembly of the two largest subunits of Pol II^43^. RPAP2 would then anchor GPN1-3 to assembled Pol II-Gdown1 and all proteins would translocate with Pol II into the nucleus. GPN1-GPN3 would then facilitate RPAP2 recycling to the nucleus by virtue of a nuclear export signal present within GPN1 (Fig. 5f). Further work will be required to elucidate the proteins, which recognize GPN1-GPN3 in the GTP- and GDP-bound states and clarify the role of the GPN1-GPN3 complex as a chaperone.

## Methods

### Endogenous protein tagging

Wild-type K562 cells (DSMZ) were edited to express an eGFP–3C–Gdown1 fusion protein (final construct Blasticidin - P2A - EGFP-GS linker - AID degron - GS linker - 3C protease site - Gdown1) using a modified CRISPR–Cas9 knock-in protocol as previously described[ref:]. In brief, the gRNA (5’-AGAATGTGCTCGCTGCCCCG-3’) was inserted into the pLCG plasmid and 500 bp regions flanking the Gdown1 start codon from K562 genomic DNA were cloned into a pLPG. K562 cells were co-transfected with the HDR donor and Cas9 plasmids and subjected to blasticidin selection (10 μg/mL, Invitrogen). GFP-positive cells were isolated by FACS using a BD FACSAria III (BD Biosciences) and genotyped with primers GCCAGAGTCTCCCAAATCCT and ACAAGTGTCTCGTGGGAAGT. Expression of GFP–Gdown1 was verified by western blot using anti-POLR2M (HPA068141, Sigma Aldrich) and anti-GFP (A11122, Invitrogen) antibodies.

### Cell fractionation

For extract preparation, human K562 cells endogenously expressing GFP-AID-3C-Gdown1 were cultured in 12 L RPMI media, 10% FBS (Gibco), 2% GlutaMax (Gibco), 1% sodium pyruvate (Sigma Aldrich) and 1% penicillin-streptomycin (Sigma Aldrich). Cytoplasmic extract (CE) and nuclear extract (NE) were prepared as previously described^44^ and dialysed against Buffer E (20 mM HEPES-KOH pH 7.9, 20% (v/v) glycerol, 100 mM KCl, 0.2 mM EDTA, 0.5 mM DTT). Nuclear and cytoplasmic separation was confirmed by Western blot using antibodies against GAPDH1 (Sigma Aldrich) and U1snRNP70 (Santa Cruz Biotechnology) as cytoplasmic and nuclear makers respectively.

### Preparation of GFP nanobody-coupled resin

A plasmid encoding a GFP-nanobody carrying a C-terminal 6xHis tag was a gift from Brett Collins (Addgene plasmid # 49172; http://n2t.net/addgene:49172; RRID:Addgene_49172)^45^. The coding sequence was altered using site-directed mutagenesis to contain two lysine residues between the C-terminus of the insert and the 6xHis tag (GFP nanobody-KK-6xHis), to improve coupling to AminoLink Plus resin. The modified GFP nanobody was expressed in E. coli BL21(DE3)RIL cells. The culture was grown to an optical density 600 nm (OD600) of 0.7 at 37 °C and then induced by 0.5 mM IPTG for 16 hrs at 18°C. Cells were collected and resuspended in NaPi buffer (50 mM sodium phosphate pH 7.4 at 25 °C, 200 mM NaCl) with addition of protease inhibitors (PIs, 1 mM PMSF, 1 mM benzamidine, 60 μM leupeptin, and 200 µM pepstatin) and 10 mM imidazole, then sonicated on ice. GFP nanobody was purified using a HisTrap column HP (Cytiva) equilibrated in NaPi buffer with PIs and 10mM imidazole. The column was washed with NaPi buffer containing 30 mM imidazole, incubated with ATP Wash buffer (NaPi buffer, 30 mM imidazole, 5 mM ATP, 2 mg/mL denatured *E. coli* protein) at 25 °C to remove chaperones and then equilibrated back to 4 °C. GFP nanobody was eluted using a linear gradient of imidazole in NaPi buffer (30 mM to 500 mM). Peak protein fractions were subjected to size exclusion chromatography using a Superdex Increase 75 10/300 GL (Cytiva) equilibrated in Coupling buffer (50 mM sodium phosphate pH 7.2 at 25 °C, 150 mM NaCl). Purified GFP nanobody was concentrated using 10,000 MWCO Amicon Ultra Centrifugal Filter (Merck Millipore) and stored at −80 °C until use.

The GFP-nanobody resin was prepared using purified GFP nanobody coupled to AminoLink Plus resin (0.3mg/100 uL resin, Thermo Scientific) according to the manufacture’s protocol and stored in 1x PBS supplemented with 0.02% sodium azide at 4°C.

### Purification of human Gdown1-associated proteins

The GFP-nanobody resin (homemade for biochemical and structural experiments, GFPtrap (Chromotek) for western blot and mass spectrometry experiments) was first equilibrated using Buffer A (20 mM HEPES-NaOH pH 7.9 at 25°C, 100 mM NaCl, 0.05% Triton X-100, 2 mM MgCl_2_, 10% (v/v) glycerol, 2 mM DTT). CE or NE was then applied to the resin after dilution 1:1 with Dilution buffer (20 mM HEPES-KOH pH 7.9 at 25°C, 7.5 mM KCl, 3 mM MgCl_2_, 1 mM DTT) and incubated for 2 hours at 4 °C. The resin with bound proteins was washed Buffer A and once with Buffer B (20 mM HEPES-NaOH (pH 7.9 at 25°C), 100 mM NaCl, 2 mM MgCl_2_, and 2 mM DTT). Protein elution was performed using 3C protease for 2 hours at 4 °C. Proteins intended for mass spectrometry analysis were flash-frozen in liquid nitrogen and stored at −80 °C. Cytoplasmic RNA polymerase II complexes destined for XL-MS or cryo-EM were subjected to additional processing as detailed below.

### Mass spectrometry analysis of Gdown1-associated proteins

Digestion, reduction, and alkylation were carried out as previously described^46^ (project ID: PXD069399). The nano HPLC system (UltiMate 3000 RSLC nano system, Thermo Fisher Scientific) was coupled to an Q Exactive HF-X mass spectrometer equipped with a Proxeon nanospray source. Peptides were loaded onto a trap column (PepMap Ac-claim C18, 5 mm × 300 μm ID, 5 μm particles, 100 Å pore size, Thermo Fisher Scientific) at a flow rate of 25 μl/min using 0.1% trifluoroacetic acid (TFA) as mobile phase. Peptides were analysed on an analytical column (PepMap Acclaim C18, 500 mm × 75 μm ID, 2 μm, 100 Å, Thermo Fisher Scientific) with a linear gradient from 2-35% B (80% acetonitrile (ACN), 0.1% formic acid (FA)). The Q Exactive HF-X mass spectrometer was operated in data-dependent mode, performing a full scan (m/z range 380-1500, resolution 60,000), followed each by MS/MS scans of the most abundant ions.

MS/MS were acquired with an isolation window of 1.0 m/z and fragmentation was achieved with HCD with a collision energy of 28% NCE. For peptide identification, the RAW-files were loaded into Proteome Discoverer (version 2.3.0.523, Thermo Scientific). All MS/MS spectra were searched using MSAmanda v2.0.0.12368^47^. For the protein-identification-set, the peptide mass tolerance was set to ±5 ppm and fragment mass tolerance to ±8 ppm, the maximum number of missed cleavages was set to 2, using tryptic enzymatic specificity with proline restriction. Peptide and protein identification was performed in two steps. First, the RAW-files were searched against the human_uniprot_reference_2020-12-05.fasta with beta-methylthiolation on cysteines as fixed modification. The result was filtered to 1 % FDR on protein level using the Percolator algorithm^48^. Proteins were filtered to be identified by a minimum of 2 PSMs in 1 sample. Peptides were subjected to label-free quantification using IMP-apQuant^49^. Proteins were quantified by summing unique and razor peptides and applying intensity-based absolute quantification (iBAQ, Schwan-häusser et al., 2011). Protein-abundance-normalization was done by sum normalization of the MaxLFQ algorithm^50^. Identified proteins were filtered to a minimum of 2 peptides. Statistical significance of differentially expressed proteins was determined by limma^51^.

### Cryo-electron microscopy of cytoplasmic Pol II complexes

Cytoplasmic Pol II complexes, recovered using GFP-nanobody affinity resin as described above, were purified using gradient fixation (GraFix)^52^. Briefly, sucrose gradients were prepared using stepwise layering. Light sucrose solution (5% sucrose, 20 mM HEPES-NaOH pH 7.9 at 25°C, 100 mM NaCl, 2 mM MgCl_2_, 2 mM DTT) and heavy sucrose solution (30% sucrose, 20 mM HEPES-NaOH pH 7.9 at 25°C, 100 mM NaCl, 0.05% glutaraldehyde) were prepared and gradients were assembled by adding successive layers containing 30%, 23.75%, 17.5%, 11.25% and 5% sucrose (w/v). Each layer was flash-frozen on dry ice for 5 minutes prior to the addition of the next. Once all layers were in place, the gradients were allowed to thaw slowly overnight at 4 °C. Gradient preparation used for separation of cytoplasmic Pol II for further mass-spectrometry band identification did not contain glutaraldehyde. Centrifugation was performed at 32,000 rpm for 16 hours at 4°C. Gradients were fractionated, and the fractions containing Pol II complexes were pooled. To quench residual crosslinker, 50 mM L-lysine (pH 7.8 at 25 °C) was added at 25°C. Samples were then dialyzed for 24 hours at 4 °C in buffer containing 20 mM HEPES-NaOH pH 7.9 at 25°C, 100 mM NaCl, 2 mM MgCl_2_, and 2 mM DTT.

Final concentration of the dialyzed complexes was achieved using 100 kDa MWCO Amicon Ultra centrifugal filters (Merck Millipore). Cytoplasmic Pol II complexes were then applied to wells and overlaid with a 2 nm carbon film, followed by a 30-minute incubation at 4°C. Quantifoil Cu 300 R1.2/1.3 grids were then floated onto the sample surface to adsorb the carbon layer. Afterwards, grids were vitrified in liquid ethane using a Vitrobot Mark IV (Thermo Scientific, 100% humidity, 4°C). Cryo-EM grids were loaded into a Thermo Fisher Titan Krios G3i transmission electron microscope (TEM) operating at 300 kV and equipped with a Gatan K3 BioQuantum direct electron detector, using an energy filter with a 10 eV slit width. Data acquisition was performed with EPU software (version 2.11). Images were captured at a nominal magnification of 130,000×, corresponding to a physical pixel size of 0.656 Å. After two collection sessions, a total of 43,150 micrographs were acquired with a defocus range of –0.3 to –3.5 μm. The electron exposure rate was approximately 40.4 e-/Å²/s, with each exposure lasting 1.86 seconds, yielding a cumulative dose of 75 e-/Å² across 50 frames, and with an exposure rate on the detector of ∼15-17 e-/pix/s.

Micrographs were pre-processed (motion correction and CTF estimation) and binned 2-fold to a final pixel size of 1.312 Å using Warp (version 1.0.9)^53^. Particle coordinates were selected using Warp, and dose-weighted particles were extracted and imported into RELION 3.1 for further processing. Data from two collection sessions were independently cleaned using a combination of 2D and 3D classification in RELION 3.1 (Extended Data Fig. 2). The two data sets were merged and subjected to CTF refinement (per-particle defocus and per-micrograph astigmatism) beam tilt refinement, and particle polishing, before a second round of defocus and astigmatism fitting. The resulting refined density was used to generate the model of the Pol II core (excluding the stalk and clamp domains), whereas 3D classification in RELION 3.1 and RELION 5.0 was utilized to improve densities for the modeling of RPAP2 and Gdown1 (Table 1, Extended Data Fig. 2). Local resolution was calculated in cisTEM (2.0.0 alpha) using the non-fixed resolution estimator described in Rohou 2020^54^. ChimeraX (version 1.10)^55^ was used for visualization.

**Table 1.**
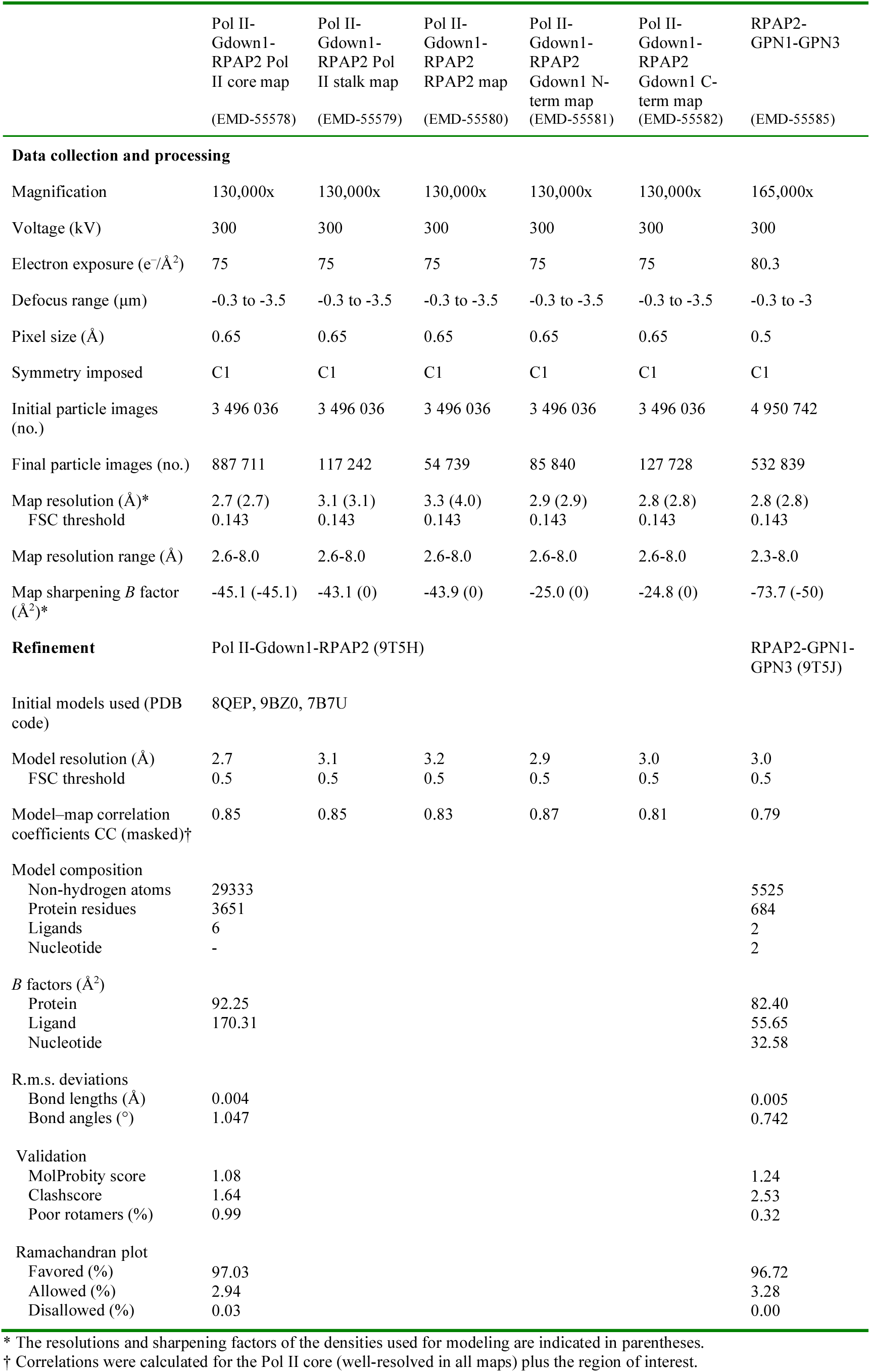
Cryo-EM data collection, refinement, and validation statistics.

|  | Pol II-Gdown1-RPAP2 Pol<br>II core map<br>(EMD-55578) | Pol II-Gdown1-RPAP2 Pol<br>II stalk map<br>(EMD-55579) | Pol II-Gdown1-RPAP2<br>RPAP2 map<br>(EMD-55580) | Pol II-Gdown1-RPAP2<br>Gdown1 N-term map<br>(EMD-55581) | Pol II-Gdown1-RPAP2<br>Gdown1 C-term map<br>(EMD-55582) | RPAP2-GPN1-GPN3<br>(EMD-55585) |
| --- | --- | --- | --- | --- | --- | --- |
| **Data collection and processing** |  |  |  |  |  |  |
| Magnification | 130,000x | 130,000x | 130,000x | 130,000x | 130,000x | 165,000x |
| Voltage (kV) | 300 | 300 | 300 | 300 | 300 | 300 |
| Electron exposure (e<sup>-</sup>/Å<sup>2</sup>) | 75 | 75 | 75 | 75 | 75 | 80.3 |
| Defocus range (μm) | -0.3 to -3.5 | -0.3 to -3.5 | -0.3 to -3.5 | -0.3 to -3.5 | -0.3 to -3.5 | -0.3 to -3 |
| Pixel size (Å) | 0.65 | 0.65 | 0.65 | 0.65 | 0.65 | 0.5 |
| Symmetry imposed | C1 | C1 | C1 | C1 | C1 | C1 |
| Initial particle images (no.) | 3 496 036 | 3 496 036 | 3 496 036 | 3 496 036 | 3 496 036 | 4 950 742 |
| Final particle images (no.) | 887 711 | 117 242 | 54 739 | 85 840 | 127 728 | 532 839 |
| Map resolution (Å)* | 2.7 (2.7) | 3.1 (3.1) | 3.3 (4.0) | 2.9 (2.9) | 2.8 (2.8) | 2.8 (2.8) |
| FSC threshold | 0.143 | 0.143 | 0.143 | 0.143 | 0.143 | 0.143 |
| Map resolution range (Å) | 2.6-8.0 | 2.6-8.0 | 2.6-8.0 | 2.6-8.0 | 2.6-8.0 | 2.3-8.0 |
| Map sharpening *B* factor (Å<sup>2</sup>)* | -45.1 (-45.1) | -43.1 (0) | -43.9 (0) | -25.0 (0) | -24.8 (0) | -73.7 (-50) |
| **Refinement** | Pol II-Gdown1-RPAP2 (9T5H) |  |  |  |  | RPAP2-GPN1-GPN3 (9T5J) |
| Initial models used (PDB code) | 8QEP, 9BZ0, 7B7U |  |  |  |  |  |
| Model resolution (Å) | 2.7 | 3.1 | 3.2 | 2.9 | 3.0 | 3.0 |
| FSC threshold | 0.5 | 0.5 | 0.5 | 0.5 | 0.5 | 0.5 |
| Model-map correlation coefficients CC (masked)† | 0.85 | 0.85 | 0.83 | 0.87 | 0.81 | 0.79 |
| Model composition |  |  |  |  |  |  |
| Non-hydrogen atoms | 29333 |  |  |  |  | 5525 |
| Protein residues | 3651 |  |  |  |  | 684 |
| Ligands | 6 |  |  |  |  | 2 |
| Nucleotide | - |  |  |  |  | 2 |
| *B* factors (Å<sup>2</sup>) |  |  |  |  |  |  |
| Protein | 92.25 |  |  |  |  | 82.40 |
| Ligand | 170.31 |  |  |  |  | 55.65 |
| Nucleotide |  |  |  |  |  | 32.58 |
| R.m.s. deviations |  |  |  |  |  |  |
| Bond lengths (Å) | 0.004 |  |  |  |  | 0.005 |
| Bond angles (°) | 1.047 |  |  |  |  | 0.742 |
| Validation |  |  |  |  |  |  |
| MolProbity score | 1.08 |  |  |  |  | 1.24 |
| Clashscore | 1.64 |  |  |  |  | 2.53 |
| Poor rotamers (%) | 0.99 |  |  |  |  | 0.32 |
| Ramachandran plot |  |  |  |  |  |  |
| Favored (%) | 97.03 |  |  |  |  | 96.72 |
| Allowed (%) | 2.94 |  |  |  |  | 3.28 |
| Disallowed (%) | 0.03 |  |  |  |  | 0.00 |

\* The resolutions and sharpening factors of the densities used for modeling are indicated in parentheses.
† Correlations were calculated for the Pol II core (well-resolved in all maps) plus the region of interest.

### Pol II-Gdown1-RPAP2 model building and refinement

A starting model for Pol II was generated by mutating a previously published model of porcine Pol II lacking the clamp domain (8QEP^56^) to include the corresponding human residues (T74I, S126T). A starting model for the Pol II stalk (RPB4/RPB7) was obtained from a previously published high-resolution Pol II structure (9BZ0^57^). Prediction of a Pol II (no clamp)-RPAP2 structure using AlphaFold3^58^ was combined with a previously solved Pol II-RPAP2 structure (7B7U^23^) in order to assign an identity to residues previously designated as “unknown”. An AlphaFold3 prediction of Pol II (no clamp) bound to Gdown1 was used to generate a starting model for Gdown1.

After fitting the starting model into the core density, areas of poor fit or geometry were adjusted using Isolde (version 1.10.1)^59^ and real space refined using Phenix (version 1.21.2)^60^, applying Isolde-generated restraints (global minimization with reference restraints and ADP refinement). This procedure was used to fit the RPAP2 and Gdown1 N-terminus (15-68) models into the densities. The model for the Gdown1 C-terminus (226-336) was adjusted using Isolde prior to Phenix real-space refinement.

The Pol II core was well-resolved in all classified densities, and was included in all refinements to maintain the interfaces between Pol II and RPAP2 or Gdown1. The final Pol II-Gdown1-RPAP2 model was generated by aligning the Pol II core of all refined models, then merging the fitted RPAP2 and refined Gdown1 regions into a composite model.

### Pol II-GPN1-GPN3 structure prediction

The Pol II-GPN1-GPN3 and Pol II-TTF2 (UniProt Q9UNY4) structures were predicted using the AlphaFold3 webserver. Visualization and calculation of Coulombic electrostatic potentials were carried out using UCSF ChimeraX.

### Constructs for recombinant RPAP2 and GPN1-GPN3 expression

The open reading frame of human RPAP2 (UniProt Q8IXW5) was cloned into a modified pFastBac-derived vector (438-C) containing an N-terminal 6xHis-MBP tag followed by a 3C protease site[ref: Gradia et al. 2017]. A truncated version of RPAP2, encompassing amino acids 1–215, was cloned into pOPINB *E. coli* vector with an N-terminal 6xHis and 3C protease site^61^. Human GPN1 (UniProt Q9HCN4) was cloned into a second modified pFastBac-derived vector (438-B) to generate a construct containing an N-terminal 6xHis-3C protease site, whereas GPN3 (UniProt Q9UHW5) was cloned into the vector lacking any tag. GPN1 and GPN3 were further subcloned into the same 438 vector using ligation-independent cloning as previously described^62^.

### Protein expression and purification

GPN1 and GPN3 were co-expressed in insect cells. Cells were resuspended in GPN buffer (20 mM HEPES-NaOH pH 7.9 at 25 °C, 200 mM NaCl, 1 mM DTT) with 30 mM imidazole and PIs and sonicated on ice. Cleared lysate was applied to a HisTrap HP column (Cytiva) equilibrated in GPN buffer supplemented with 30 mM imidazole and PIs. Column was washed with GPN buffer with 50 mM imidazole. GPN1-GPN3 complex was eluted with a linear gradient of 50 mM to 500 mM imidazole and cleaved with 3C protease over night while dialyzing against GPN buffer and 30 mM imidazole. Cleaved proteins were applied to a HisTrap column and flow-through collected. Proteins were separated on a Superdex Increase 75 10/300 GL column (Cytiva) equilibrated in GPN size exclusion buffer (20 mM HEPES pH 7.9 at 25 °C, 100 mM NaCl, 2 mM MgCl_2_, 1 mM DTT). Peak fractions were pooled. For purification of His-GPN1-GPN3 complex for pulldown experiments, protease cleavage and reverse Ni-NTA steps were omitted.

Full-length RPAP2 was expressed in insect cells as previously described^23^. The RPAP2 Pol II-binding domain (RPAP2 (1-215)) was expressed in E. coli BL21(DE3)RIL cells. Cells were grown in LB with 30 µg/mL kanamycin and 100µg/mL chloramphenicol to OD600 0.7 and protein expressed adding 0.5 mM IPTG for 16 hours at 18 °C. RPAP2 full-length and RPAP2 (1-215) protein were purified as described previously with small changes^23^. Briefly, cells were lysed by sonication in RPAP2 buffer (25 mM Tris-HCl pH 8.0 at 25°C, 500 mM NaCl, 30 mM imidazole, 10% (v/v) glycerol, 10 µM ZnCl_2_, 1 mM DTT) with addition of PIs. The cleared lysates were applied to a HisTrap HP column (Cytiva) equilibrated in RPAP2 buffer with PIs. Two MBPtrap columns (Cytiva) equilibrated in RPAP2 buffer were connected in tandem to the HisTrap column and bound proteins were eluted from HisTrap column using linear gradient of RPAP2 buffer from 30 mM to 500 mM imidazole. Afterwards, MBPTrap columns were washed with RPAP2 buffer and elution performed in RPAP2 buffer containing 100 mM maltose. RPAP2 fractions were pooled and cleaved with 3C protease over night at 4°C and re-applied on HisTrap column equilibrated in RPAP2 buffer. Flow-through was collected and applied on a Superdex Increase 200 10/300 column (Cytiva) equilibrated in RPAP2 size exclusion buffer (20 mM HEPES-NaOH pH 7.9 at 25 °C, 200 mM NaCl, 10% (v/v) glycerol, 10 µM ZnCl_2_, 1 mM DTT). Fractions containing RPAP2 were pooled and concentrated using a 30,000 MWCO Amicon Ultra Centrifugal Filter (Merck Millipore). RPAP2 (1-215) was purified using the same procedure, omitting the MBPtrap column step.

Porcine Pol II and human Gdown1 were purified as previously described^63^, except that for Gdown1 expression, the culture was grown to OD600 0.7 at 37°C before induction of expression by addition of 1 mM IPTG for 6 hrs at 25°C. To prepare Pol II-Gdown1 complex, Pol II was incubated with human Gdown1 in a 1:3 molar ratio and incubated at 4 °C for 30 minutes, prior to purification over a Sephacryl S-300 HiLoad column equilibrated in Pol II buffer (150 mM NaCl, 5 mM HEPES-KOH pH 7.25 at 25 °C, 10 μM ZnCl_2_, 10 mM DTT). Peak fractions were concentrated as described above. All proteins were aliquoted and frozen in liquid nitrogen prior to storage at −80 °C.

### GTPase activity assay

GTPase activity was measured by using Malachite Green Phosphate Assay kit (Sigma-Aldrich) following the manufacturer instructions. 0.66 µM GPN1-GPN3 were incubated with 0.25 mM GTP (Thermo Fischer Scientific) for 60 min at 37°C in GPN size exclusion buffer.

### Pull-down interaction assays

His-GPN1-GPN3 and Pol II or Pol II EC were mixed in 1:2 molar ratio in Low salt buffer (20 mM HEPES pH 7.9 at 25 °C, 100 mM NaCl, 2 mM MgCl_2_, 1 mM DTT) and incubated for 30 minutes at 4°C and then applied on MagneHis beads (Promega) for 30 minutes at 4°C. Unbound proteins were washed away with High salt wash buffer (20 mM HEPES pH 7.9 at 25°C, 300 mM NaCl, 2 mM MgCl_2_, 1 mM DTT). 0.66 µM HisGPN1-GPN3 was incubated with 10 mM GTP (Thermo Scientific), 10 mM GMPPNP (Jena Bioscience) or 10 mM GDP (Jena Bioscience) for 30 minutes at 37 °C and washed once with Low salt buffer. 0.72 µM RPAP2 or 1.3 µM RPAP2(1-215) with addition of 10 mM GTP, 10 mM GMPPNP or 10 mM GDP was added to the beads and incubated for 30 minutes at 4°C. Beads were washed with High salt wash buffer supplemented with 30 mM imidazole. Immobilized proteins were eluted with Low salt buffer supplemented with 500 mM imidazole. The proteins were analyzed by SDS-PAGE gel stained with Coomassie.

### Mass photometry

For Pol II complex formation, Pol II and Pol II-Gdown1 complex with GPN1-GPN3 and/or RPAP2 was prepared by incubating the proteins in 1:2:2 molar ratio for 30 minutes at 4 °C. Samples were diluted to a final concentration of 100 nM using Dilution buffer (20 mM HEPES-NaOH pH 7.9 at 25 °C, 100 mM NaCl, 2 mM MgCl_2_, 10 µM ZnCl_2_, 1 mM DTT). Samples and respective controls were measured using Refeyn Two MP mass photometer (Refeyn Ltd.) at a final concentration 20 nM. The movies were collected for 60 seconds. The movies were processed with DiscoverMP software. The instrument was calibrated using BSA or MfP1 standard (Refeyn Ltd.) before each measurement.

For RPAP2-GPN1-GPN3 assembly experiments performed without added nucleotide, GPN1-GPN3 and RPAP2 were added in a 1:1 molar ratio and incubated for 30 minutes at 4 °C prior to measurement. For nucleotide mass-photometry experiments, GPN1-GPN3 was incubated with 10 mM GTP (Thermo Scientific), 10 mM GMPPNP (Jena Bioscience) or 10 mM GDP (Jena Bioscience) for 30 minutes at 37 °C and then RPAP2 was added in a 1:1 molar ratio and incubated for 30 minutes on ice.

### Chemical crosslinking and mass spectrometry

Cytoplasmic Pol II samples were enriched using ProteoMiner protein enrichment kit (Bio-Rad Laboratories). For crosslinking, 50 µL of disuccinimidyl sulfoxide (DSSO, Cayman Chemical) was added at a final concentration of 1.5 mM and incubated for 6 minutes and quenched with three µL of 1 M ammonium bicarbonate (Sigma-Aldrich). For the in vitro reconstituted sample, Pol II, Gdown1, RPAP2 and GPN1-GPN3 were mixed in a molar ratio of 1:3:2:2 and incubated for 30 minutes at 4°C. The complex was incubated for 30 minutes with 3 mM BS3 (Thermo Fisher) at 4°C and quenched with a final concentration of 60 mM ammonium bicarbonate (Sigma-Aldrich).

The crosslinked proteins were denatured by adding 8 M urea and reduced and alkylated using TCEP and IAA respectively. Samples were digested using Lys-C (FUJIFILM Wako Chemicals U.S.A. Corporation) for 1h and after dilution to 1 M urea with trypsin (Promega) overnight. Afterwards samples were acidified with TFA, peptides were cleaned using Sep-Pak tC18 1 cc Vac Cartridges (Waters) and dried in a vacuum centrifugation. Peptides were dissolved in 20 µL SEC buffer (77.5 % H2O, 22.5 % ACN, 0.1 % TFA) and loaded onto a Superose 30 Increase column (Cytiva) to enrich fractions containing crosslinked peptides^64^. Samples containing crosslinked peptides were dried and dissolved in mobile phase A (98% H2O, 2% ACN, 0.1% FA) and injected onto an Ultimate 3000 RSLCnano System coupled to an Orbitrap Eclipse Tribrid Mass Spectrometer (both Thermo Fisher Scientific). Peptides were separated on a PepMap RSLC EASY-Spray column (C18, 2 µm, 100 Å, 75 µm x 15 cm, Thermo Fisher Scientific). Separation occurred at 300 nL·min-1 with a linear gradient from 2-35% mobile phase B (2% H2O, 98% ACN, 0.1% FA) within 60 min resulting in a total method time of 94 min. Orbitrap Eclipse was equipped with FAIMS Pro alternating between three different CVs (−50,−60,−75) operating in positive ionization mode. MS1 scans were recorded at a resolution of 60,000 @200 m/z in a range from 400-1600 m/z. Precursors with a charge of 3-8 were isolated with a 1.6 m/z window and fragmented with HCD with a stepped collision energy of 21,27 and 33 % NCE. MS2 spectra were recorded at a resolution of 30,000 @200 m/z.

DSSO crosslinked samples were searched using MS Annika^65^ within the Proteome Discoverer software suite against the same database as the BS3 samples. The predefined DSSO workflow was used and CSM and crosslink level FDR was set to 1 % and only high confidence identifications were considered for the subsequent analysis. Spectra were manually verified using the PD inbuilt spectra viewer. For BS3, raw files were converted to mgf using MSConvert^66^ and searched using xiSEARCH (v1.8.5^18^). Database consisted of the expected polymerase subunits and 40 additional proteins representing the most abundant proteins identified in a corresponding proteome profiling. To evaluate the identified crosslinks xiFDR (v2.3) was employed at a maximum Residue Pair FDR of 1% crosslink and identifications were also manually verified using xiView^67^.

### Cryo-electron microscopy of reconstituted RPAP2-GPN1-GPN3 complex

GPN1-GPN3 and RPAP2 were incubated in a 1:1 molar ratio for 30 minutes at 4°C. Samples were subjected to size exclusion on a Superdex 200 Increase 3.2/300 column (Cytiva) equilibrated in buffer (cooled to 4°C) containing 20 mM HEPES-NaOH pH 7.9 at 25°C, 100 mM NaCl, 2 mM MgCl_2_, 10 µM ZnCl_2_, 1 mM DTT.

Fractions containing the RPAP2-GPN1-GPN3 complex were pooled and concentrated to 0.6 mg/mL using a 30,000 MWCO Amicon Ultra Centrifugal Filter (Merck Millipore). 3.5 µL of the sample was applied to Quantifoil R0.6/1 holey carbon grids that had been glow discharged for 40 sec (25 mA current, 7.0 x 10^-1^ mbar vacuum). Using a Vitrobot Mark IV set to 100% humidity and 4 °C, grids were blotted for 7 seconds and immediately plunge frozen in liquid ethane and stored in liquid nitrogen. Cryo-EM grids were imaged using a Thermo Fisher Titan Krios G3i TEM operating at 300 kV and equipped with a Gatan K3 BioQuantum direct electron detector, using an energy filter with a 10 eV slit width. Data acquisition was performed using EPU software (version 2.11). Images were captured at a nominal magnification of 165,000×, corresponding to a physical pixel size of 0.50 Å. A total of 14 362 micrographs were collected with a defocus range of –0.4 to –3 μm. The electron exposure rate was 69.4 e-/Å²/s, with each exposure lasting 1.16 seconds, yielding a cumulative dose of 80.3 e-/Å² across 80 frames. The exposure rate on the detector was ∼16 e-/pix/s.

Micrographs were pre-processed in RELION 5.0, and CTF estimation was performed using CTFFIND-4.1^68^. Particle coordinates were selected in cryoSPARC (version 4.4.1)^69^ using the Blob Picker. Extracted particles were cleaned first using cryoSPARC (2D classification, followed by heterogeneous refinement). To improve apparent anisotropy in the resolution of the resulting map, particles were re-imported to RELION 5.0 and subjected to additional 3D classification. Final refined densities were generated in RELION using blush regularization.

### GPN1-GPN3-RPAP2 model building and refinement

A model for the GPN1-GPN3-RPAP2 complex containing two molecules of GDP and two Mg^2+^ ions was generated using AlphaFold3. RPAP2 1-311 was deleted and the molecule was rigid body fit into the RPAP2-GPN1-GPN3 map, sharpened with a b-factor of −50, in ChimeraX. Regions of the model that did not agree with the density were deleted, and the model was manually rebuilt and adjusted in Coot^70^ and using Isolde. Unassigned density around GPN1-GPN3 was assigned to RPAP2 based on side chain densities observed in the cryo-EM map, with RPAP2-GPN1 crosslinks used for validation. Phenix real space refinement was then carried out using Isolde-suggested parameters (global minimization with reference restraints and ADP refinement). A DeepEMhancer-postprocessed map was used for display only^71^.

## Acknowledgements

We thank A. Salmazo for assistance with Pol II purification. We thank staff at the VBCF Proteomics facility for immunoprecipitation-mass spectrometry analysis, and J.A. Stopp for assistance with IP-MS data visualization. This research was further supported by the Scientific Service Units (SSUs) of IST Austria through resources provided by the Lab Support Facility (LSF), Electron Microscopy Facility (EMF), Scientific Computing (SciComp), and the Preclinical Facility (PCF).

## Author contributions

A.H. performed experiments and analyzed the data, unless otherwise described. B.N. performed crosslinking-mass spectrometry analysis, supervised by F.H.. U.S. generated the Gdown1 (POLR2M) GFP-tagged CRISPR K562 cell line, supervised by C.P.. A.H. and C.B. processed cryo-EM data. C.B. designed and supervised research. A.H. and C.B. prepared the manuscript, with input from all authors.

## Competing interests

The authors declare no competing interests.

## Extended Data Figures

**Extended Data Figure 1.**
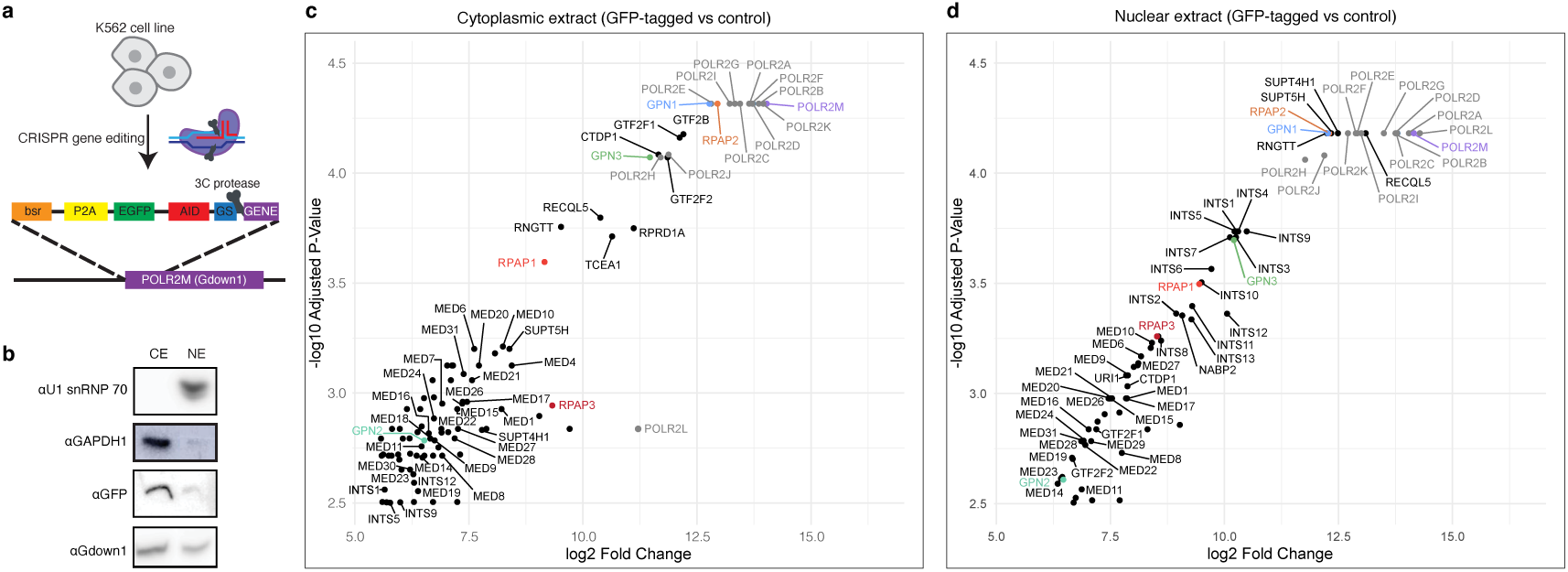
Isolation and compositional analysis of endogenous Gdown1-associated complexes. **a**, Schematic of the edited Gdown1 endogenous locus in the CRISPR-edited K562 cell line. bsr –blasticidin resistance marker; P2A-intervening porcine teschovirus-1 2A; AID - auxin-inducible degron domain; GS - Gly-Ser linker. **b**, Western blot analysis of representative cytoplasmic and nucleoplasmic extract preparations, probing for the cytoplasmic marker (GAPDH), nuclear marker (U1snRNP70). Expression of Gdown1 confirmed by probing against GFP and Gdown1. **c**, Expanded zoom-in view of the Volcano plot of cytoplasmic Gdown1-enriched factors shown in Fig. 1b. Plot generated in R-studio. **d**, Expanded zoom-in view of the Volcano plot of nucleoplasmic Gdown1-enriched factors shown in Fig. 1c. Plot generated in R-studio. Pol II subunits, gray; GPN1-3, blue/green; RPAP1-2, orange/red; Gdown1 (POLR2M), purple.

**Extended Data Figure 2.**
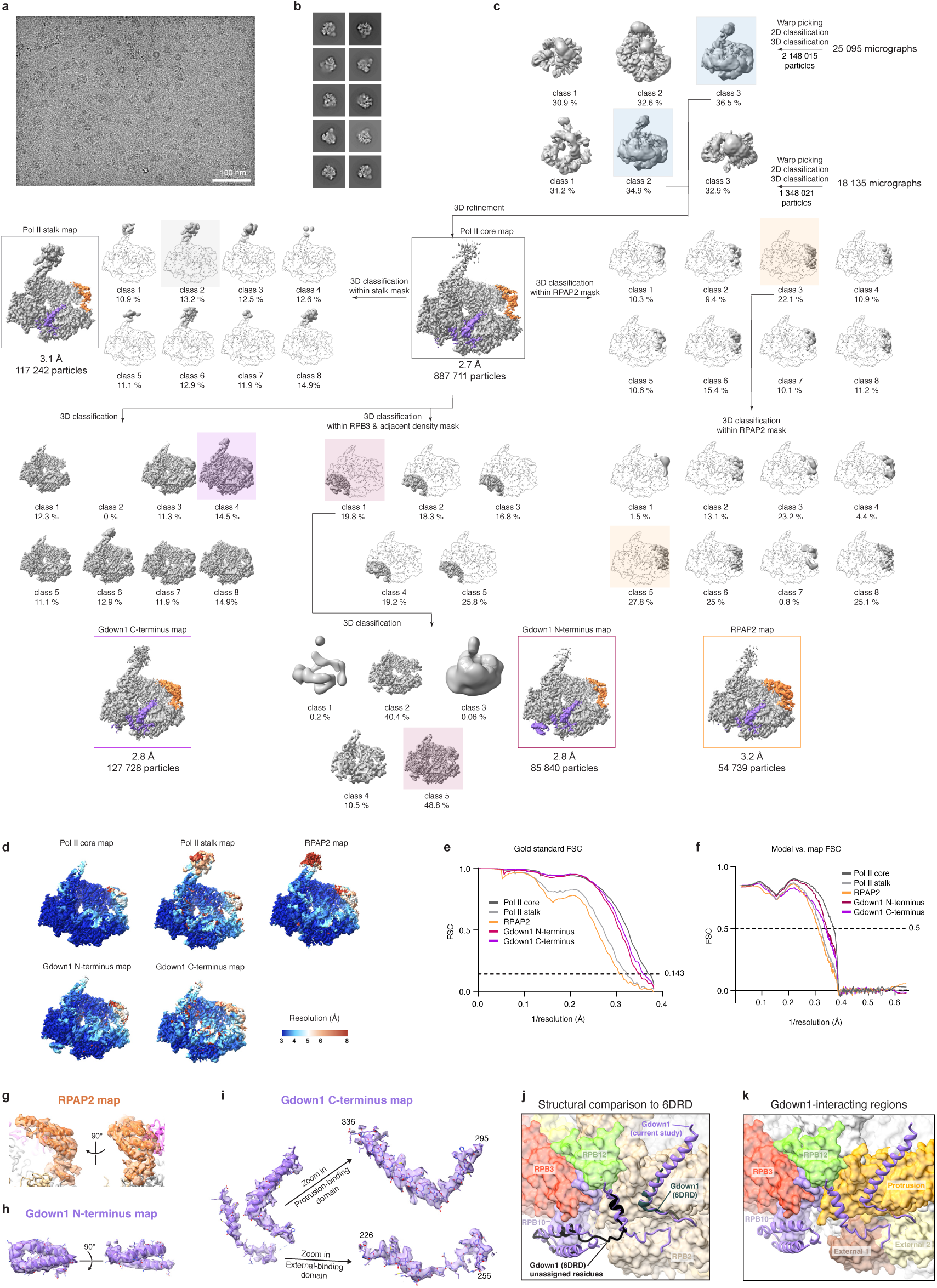
Cryo-EM analysis of cytoplasmic Pol II. **a**, Representative cryo-EM micrograph field of view at −1.3 μm defocus. Scale bar 100 nm. **b**, Representative 2D classes **c**, Processing tree outlining the image processing pipeline used to generate the five final cryo-EM densities. **d**, Non-b-factor sharpened maps colored by estimated local resolution. **e**, Fourier shell correlation plots for the five final cryo-EM densities. **f**, Model-versus-map correlation for the five final reconstitutions. **g**, Cryo-EM density of RPAP2 with the model superimposed. **h**, Cryo-EM density of the Gdown1 N-terminus (RPB10-binding domain) with the model superimposed. **i**, Left, the Gdown1 C-terminus model overlayed with cryo-EM density. Right, zoom-ins of the protrusion- and external-binding domains of Gdown1. **j**, Comparison of the Pol II-Gdown1-RPAP2 structure with a previously published Pol II-Gdown1 structure (PDB ID 6DRD^15^). Coloring is as follows: RPB2, tan; RPB3, red; RPB10, violet; RPB12, green; Gdown1 (this study, ribbon), purple; Gdown1 (PDB ID 6DRD, ribbon): assigned residues, slate; unassigned residues, black. **k**, Regions of Pol II that interact with Gdown1. RPB2 protrusion, gold; RPB2 external 1, brown; RPB2 external 2, light yellow; RPB3, red; RPB10, violet; RPB12, green; Gdown1 (ribbon), purple.

**Extended Data Figure 3.**
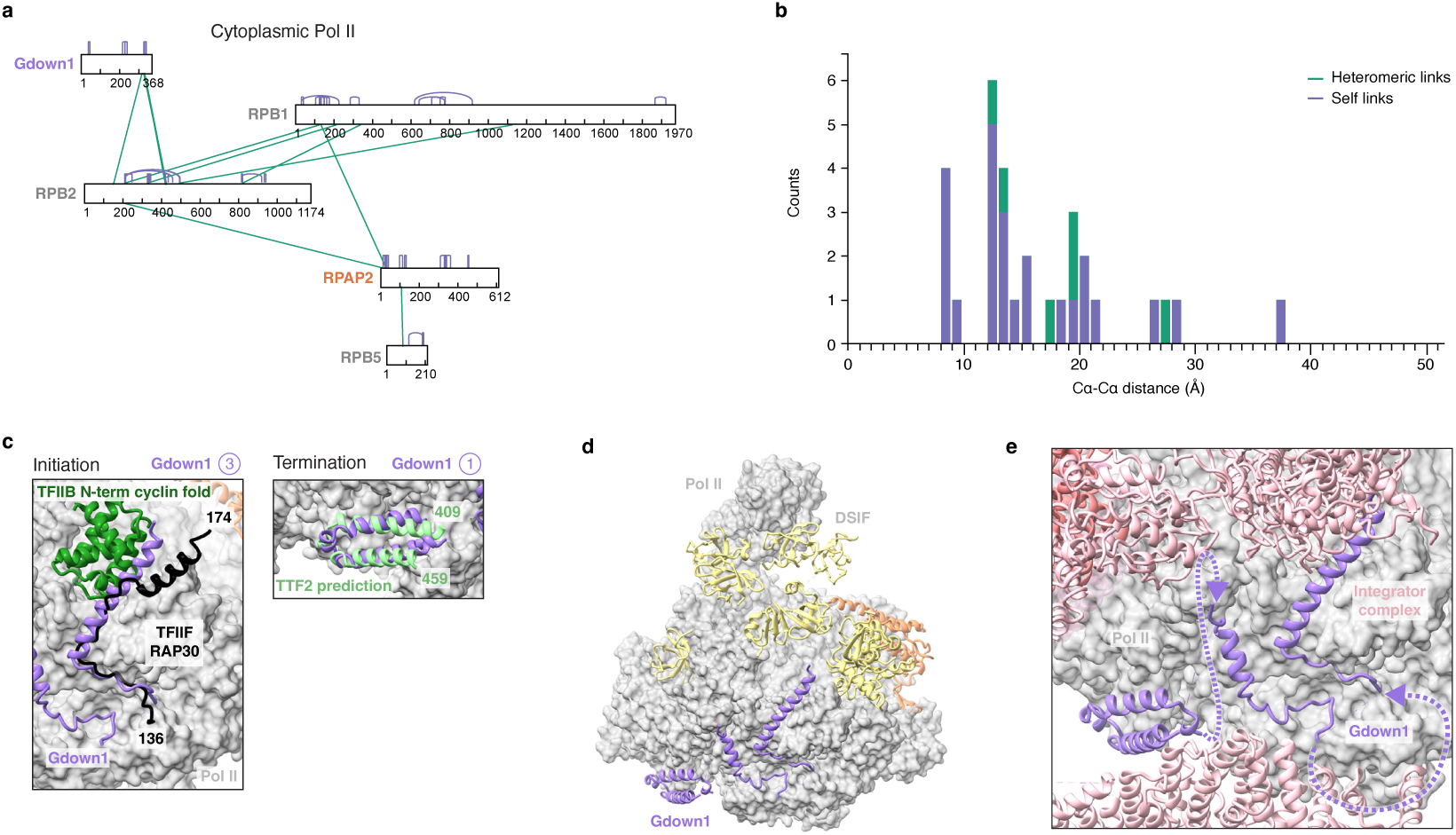
Architecture of cytoplasmic Pol II and comparison to other Pol II-bound structures. **a**, Diagram of identified crosslinks in the cytoplasmic Pol II. Violet, self crosslinked peptides; green, heteromeric crosslinked peptides. **b**, Cα-Cα distance of identified crosslinked peptides of cytoplasmic Pol II samples mapped to the Pol II-Gdown1-RPAP2 structure. Generated by xiView. **c**, Left, superposition of the Pol II-Gdown1-RPAP2 structure with a structure of the TFIID- and Mediator-containing preinitiation complex. The Gdown1 protrusion-binding domain, TFIIB B-core N-terminal cyclin fold, and TFIIF small subunit protrusion-binding region are shown (left, PDB ID 7ENA^31^). Right, superposition of the Pol II-Gdown1-RPAP2 structure with a Pol II-TTF2 structural prediction, showing the TTF2 region overlapping with the Gdown1 RPB10-binding domain. Colored as follows: Pol II, gray; Gdown1, purple; TFIIF, black; TFIIB, dark green; TTF2, light green. **d**, Superposition of cytoplasmic Pol II with DSIF (PDB ID 8UHD^72^). RPAP2, orange; DSIF, yellow; other coloring as in (c). **e,** Zoom-in comparison of the Pol II binding interfaces the Integrator complex (PDB 8RBX^28^) with those of Gdown1, as in Fig. 2d. The connectivity of unresolved regions of Gdown1 are indicated with purple dashed lines. Colored as follows: Integrator, pink; Integrator cleavage module, dark pink.

**Extended Data Figure 4.**
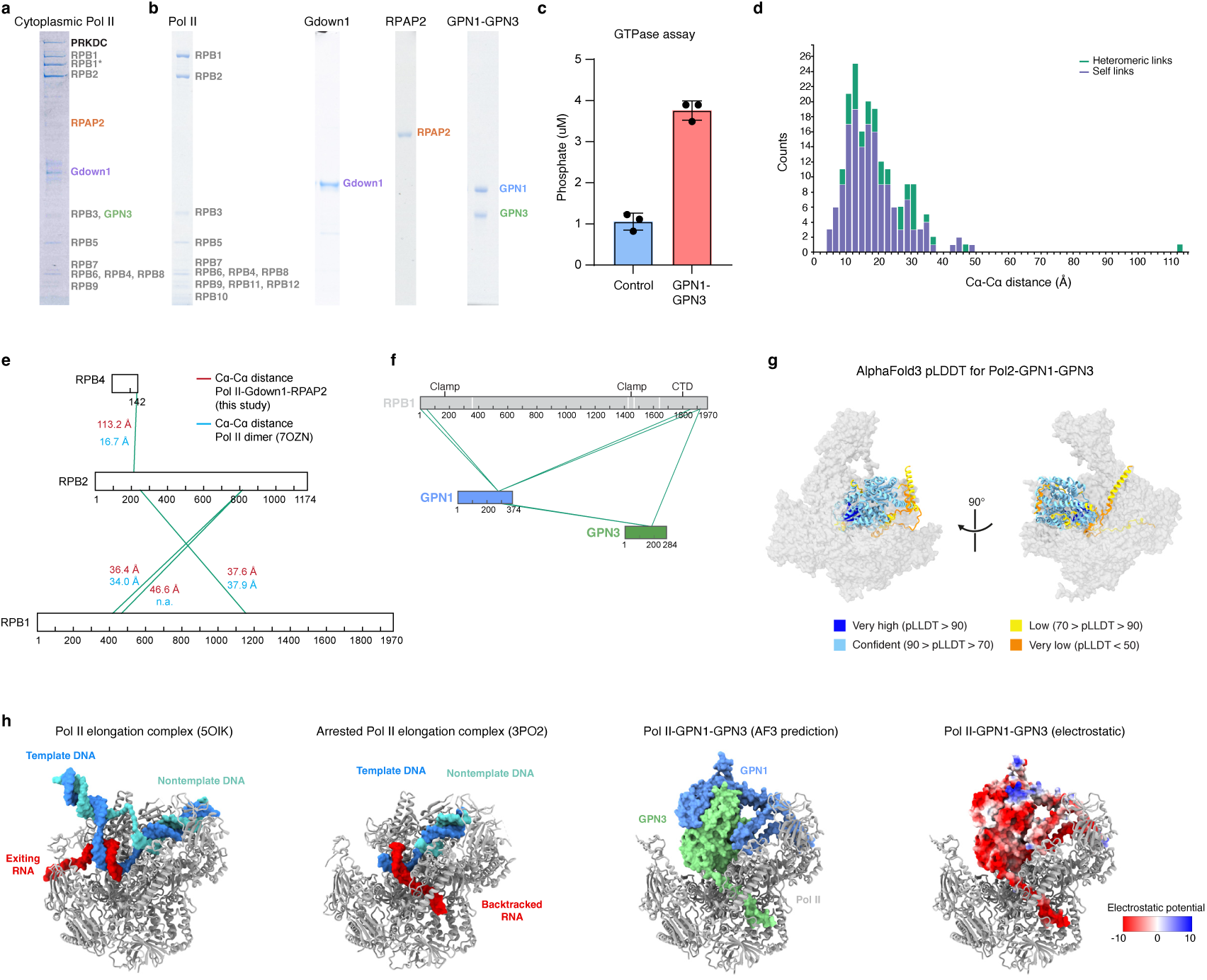
Reconstitution and characterization of Pol II-Gdown1-RPAP2-GPN1-GPN3 complexes. **a**, Representative fraction of cytoplasmic Pol II after sucrose gradient purification. Proteins identified in mass-spectrometry of excised bands are labelled. RPB1* indicates a degraded form of RPB1 with the flexible C-terminal repeat domain removed (IIb form). **b**, SDS-PAGE gels showing purified Pol II, Gdown1, RPAP2 and GPN1-GPN3 complex. **c**, GTPase activity of purified GPN1-GPN3 complex. Measured released phosphate. **d**, Cα-Cα distance of identified crosslinked peptides of in vitro Pol II-Gdown1-RPAP2-GPN1-GPN3 mapped to cytoplasmic Pol II structure. Generated by xiView^67^. **e**, Comparison of Cα-Cα distances mapped to the Pol II-Gdown1-RPAP2 structure (this study) and the Pol II dimer (PDB ID 7OZN^33^), showing the one extreme outlier that is well explained by the dimer. **f**, Diagram of identified crosslinks in the in vitro Pol II-Gdown1-RPAP2-GPN1-GPN3 sample between the RPB1 subunit and the GPN1-GPN3 complex. Violet, self crosslinked peptides; green, heteromeric crosslinked peptides. **g**, Pol II-GPN1-GPN3 structure prediction. Pol II in gray, GPN1-GPN3 prediction colored by AlphaFold3 pLLDT confidence scores. **h**, Comparison of side views of the Pol II elongation complex (PDB ID 5OIK) and arrested elongation complex (PDB ID 3PO2^35^) structures to the Pol II-GPN1-GPN3 structure prediction. Nucleic acids and GPN1-GPN3 are shown in surface representation. Far right, the same view of the Pol II-GPN1-GPN3 structure prediction, with the GPN1-GPN3 surface colored by electrostatic potential. RPB2 was removed from view for clarity.

**Extended Data Figure 5.**
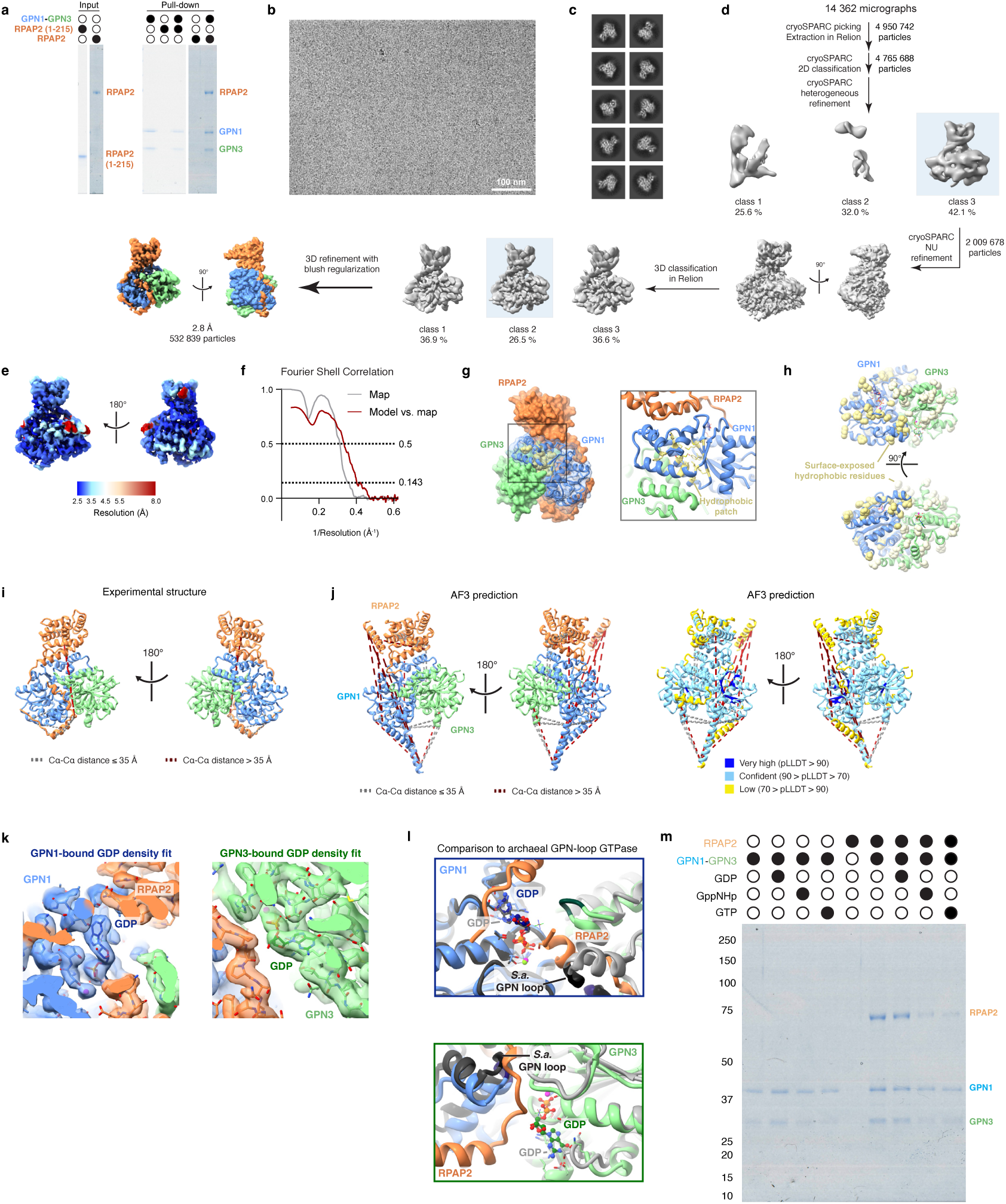
Cryo-EM and biochemical analysis of the RPAP2-GPN1-GPN3 complex. **a**, Pull-down of RPAP2 and RPAP2(1-215) via immobilized His-GPN1-GPN3. **b**, Representative cryo-EM micrograph field of view at −0.8 µm defocus. **c**, Representative 2D classes. **d**, Processing tree outlining the steps taken to generate the RPAP2-GPN1-GPN3 reconstruction. **e**, Non-b-factor sharpened map of RPAP2-GPN1-GPN3 colored by estimated local resolution. **f**, Fourier shell correlation plot for the final cryo-EM density, model-versus-map correlation for the final reconstruction and model. **g**, Overview and zoom-in view of a previously identified GPN1 hydrophobic patch in the context of the RPAP2-GPN1-GPN3 structure. Left, GPN1 surface shown in transparent blue overlayed on the GPN1 ribbon model. RPAP2, orange; GPN1, blue; GPN3, light green; residues of hydrophobic patch, yellow. **h**, View of the surface-exposed hydrophobic residues of GPN1-GPN3. Ala, Val, Cys, Pro, Leu, Iso, Met, Trp, and Phe residues with a solvent-accessible surface area cutoff of 25 (calculated in ChimeraX) are shown as spheres. GPN1 hydrophobic residues, yellow; GPN3 hydrophobic residues, light yellow. **i**, Identified crosslinked peptides in the in vitro Pol II-Gdown1-RPAP2-GPN1-GPN3 sample mapped to the RPAP2-GPN1-GPN3 structure. Cα-Cα distance ≤ 35 Å, gray; Cα-Cα distance > 35 Å, dark red. Generated by the XMAS Chimera plugin^75^. **j**, Identified crosslinked peptides in the in vitro Pol II-Gdown1-RPAP2-GPN1-GPN3 sample mapped to the AlphaFold3-predicted RPAP2-GPN1-GPN3 structure. A large number of outlier crosslinks are visible, and the AlphaFold3 model failed to predict the extended interactions of RPAP2 with GPN1 and the GPN1-GPN3 interface. Left, colored by subunit; Right, colored by pLLDT confidence score, Cα-Cα distances colored as in (i). Generated by xiView^67^. **k**, Overlay of the cryo-EM density and final RPAP2-GPN1-GPN3 model, zoomed in on the GPN1 and GPN3 GDP-binding pockets. **l**, Comparison of the human RPAP2-GPN1-GPN3 GDP binding pocket (this study) with that of the archaeal SaGPN dimer bound to GDP (PDB ID 7ZHK^37^), superimposed. Colored as follows: SaGPN monomer 1, dark gray; SaGPN monomer 2, gray; SaGPN-bound GDP, light gray, SaGPN-loop black; RPAP2-GPN1-GPN3 structure as in (a). **m**, Pull-down of RPAP2 via immobilized His-GPN1-GPN3 pre-treated with GTP, GppNHp, or GDP.

## Supplementary Material

### Supplementary Text 1 TFIIF association with Gdown1 purified from cytoplasmic extracts

Despite the well characterized competition of Gdown1 and RPAP2 with TFIIF for binding to Pol II, TFIIF was identified as enriched in the Gdown1 cytoplasmic pulldown. This observation suggests that either (1) TFIIF binds to Gdown1 as part of another complex, or (2) that either Gdown1 or TFIIF can be anchored to the Pol II complex through an additional protein adapter. We observed that CTDP1, also known as TFIIF-associating CTD phosphatase 1 (FCP1), is likewise particularly enriched in cytoplasmic pulldown. From previous high-throughput pairwise protein-protein interaction prediction^38^, CTDP1 has been identified as a highly confident binder of RPB7, part of the Pol II stalk, through its N-terminal region. CTDP1 is a well characterized binder of the small subunit of TFIIF through its central and C-terminal domains^76,77^. Thus, TFIIF may be bound to the Gdown1-Pol II complex via interaction with CTDP1. The interaction of CTDP1 with Pol II in the cytoplasm may help keep the RPB1 CTD of Pol II in a dephosphorylated state, in preparation for recruitment to promoters after import into the nucleus.s

## Notes

### Competing Interest Statement

The authors have declared no competing interest.

